# Structures of human PTP1B variants reveal allosteric sites to target for weight loss therapy

**DOI:** 10.1101/2024.08.05.603709

**Authors:** Aliki Perdikari, Virgil A. Woods, Ali Ebrahim, Katherine Lawler, Rebecca Bounds, Nathanael I. Singh, Tamar (Skaist) Mehlman, Blake T. Riley, Shivani Sharma, Jackson W. Morris, Julia M. Keogh, Elana Henning, Miriam Smith, I. Sadaf Farooqi, Daniel A. Keedy

**Author notes:** Address correspondence to Sadaf Farooqi at and Daniel Keedy at. These authors contributed equally to this work (Aliki Perdikari, Virgil A. Woods, Ali Ebrahim). **Author contributions:** AP, VAW, AE, KL, ISF, DAK designed research; AP, VAW, AE, KL, RB, NIS, TM, BTR, SS, JWM, JMK, EH, MS performed research; VAW, AE, NIS, TM analyzed data; AP, VAW, AE, KL, TM, BTR, ISF, DAK wrote the paper. **Competing interests:** ISF has consulted for a number of companies developing weight loss drugs including Eli Lilly, Novo Nordisk, and Rhythm Pharmaceuticals. The other authors declare no competing interests.

## Abstract

Protein Tyrosine Phosphatase 1B (PTP1B) is a negative regulator of leptin signaling whose disruption protects against diet-induced obesity in mice. We investigated whether structural characterization of human PTP1B variant proteins might reveal allosteric sites to target for weight loss therapy. To do so, we selected 12 rare variants for functional characterization from exomes from 997 people with persistent thinness and 200,000 people from UK Biobank. Seven of 12 variants impaired PTP1B function by increasing leptin-stimulated STAT3 phosphorylation in human cells. Focusing on the variants in and near the ordered catalytic domain, we ascribed structural mechanism to their functional effects using *in vitro* enzyme activity assays, room-temperature X-ray crystallography, and local hydrogen-deuterium exchange mass spectrometry (HDX-MS). By combining these complementary structural biology experiments for multiple variants, we characterize an inherent allosteric network in PTP1B that differs from previously reported allosteric inhibitor-driven mechanisms mediated by catalytic loop motions. The most functionally impactful variant sites map to highly ligandable surface sites, suggesting untapped opportunities for allosteric drug design. Overall, these studies can inform the targeted design of allosteric PTP1B inhibitors for the treatment of obesity.

**Significance:** Obesity is a growing public health concern worldwide. The human enzyme PTP1B is a validated obesity drug target, but to date, effective PTP1B inhibitors have not been developed. Allosteric drugs, which target parts of a protein distant from the active site, offer advantages for specific inhibition — but finding promising allosteric sites that bind ligands and convey allosteric signals remains challenging. To address this knowledge gap, we used human genetic studies in thin people to identify amino acid variants that diminish PTP1B function. We then used complementary structural biology methods to show that the variants exploit distinct allosteric wiring within PTP1B that includes ligand-enriched binding sites. This work demonstrates how a unique combination of genetics and biophysics can unveil promising allosteric sites in challenging drug target proteins.

## Introduction

Obesity causes substantial morbidity and mortality due to an increased risk of type 2 diabetes, cardiovascular disease, fatty liver disease, and some cancers (1). A new generation of anti-obesity medications (AOM), the Glucagon-like peptide-1 (GLP-1) / Gastric inhibitory polypeptide (GIP) / Glucagon receptor agonists, lead to 15–20% weight loss and are transforming the clinical care of people with obesity. However, a significant proportion of people cannot tolerate these medications due to adverse effects, and questions about their suitability for chronic use remain. As such, there is substantial interest in developing new AOM which are safe, effective, and well-tolerated.

Leptin signaling plays a pivotal role in the regulation of appetite and body weight and disruption of leptin or its receptor causes severe obesity in mice and humans. Protein Tyrosine Phosphatase 1B (PTP1B) is a negative regulator of leptin signaling in the hypothalamus, where it dephosphorylates the active site of leptin receptor-associated Janus Kinase 2 (JAK2) and decreases Signal Transducer and Activator of Transcription 3 (STAT3) phosphorylation and transcription of the anorectic neuropeptide, Pro-opiomelanocortin (POMC) (2). Brain-specific (3), leptin receptor-specific (4), and POMC-specific (5) deletion of *Ptp1b* results in mice that exhibit enhanced leptin sensitivity and are protected from high fat diet-induced obesity. As such there has been substantial interest in the development of PTP1B inhibitors for the treatment of obesity. However, this endeavor has proved to be challenging for a number of reasons. First, most small-molecule inhibitors target the catalytic site of PTP1B, but also bind to the highly homologous catalytic site of other PTPs including TCPTP (6, 7), which plays a crucial role in hematopoiesis (8). Second, compounds that bind effectively to the PTP1B active site tend to be charged, to mimic natural phosphotyrosine substrates, but this property often limits their cell-membrane permeability and bioavailability. Therefore, new approaches to inhibiting PTP1B safely and effectively are needed.

An alternative route to modulating the activity of enzymes such as PTP1B is allostery, the process by which a perturbation (such as inhibitor binding) at one site of the protein structure influences a distal site such as the active site to alter biological function. Previous work has established the existence of an allosteric network within the structure of PTP1B (9–17) and even provided initial allosteric inhibitor candidates that target different binding sites (9, 18–20). However, past allosteric drug candidates were discontinued due to limitations in efficacy and specificity; to date, no allosteric drugs for PTP1B have been approved for clinical use. A major remaining challenge is to identify new, more promising allosteric sites that can both convey allosteric signals and bind small-molecule ligands.

One strategy for uncovering promising new allosteric sites is to exploit amino acid changes distal from the enzyme’s active site that modulate function. Rare human variants from genetics studies offer a unique opportunity to pursue this strategy and thereby reveal allosteric weak points in proteins that can be subsequently targeted with small molecules. This approach has been demonstrated successfully for multiple therapeutic target proteins linked to obesity (21–23). Amino acid sequence changes have been leveraged to study allostery in PTP1B in the past, including coevolutionary signals from statistical coupling analysis (SCA) (11, 15, 24) and reciprocal mutations guided by nuclear magnetic resonance (NMR) chemical shift changes (13). However, to our knowledge, no studies have used human genetics to identify and characterize allosteric sites in PTP1B.

Here we report 12 rare human variants throughout the gene encoding PTP1B (*PTPN1*) that derive from a large cohort of ∼1000 thin, healthy people and participants in the UK Biobank. We demonstrate that most of these variants alter PTP1B function in human cell lines. We next focus on the variants within the ordered catalytic domain, which is capable of structural allostery (9–18, 25) and more amenable to detailed biophysical interrogation. To understand the allosteric mechanisms underlying the functional effects of these variants, we use complementary structural biology techniques that are sensitive in different ways to protein structural flexibility. First, we use room-temperature (RT) X-ray crystallography, which reveals all-atom conformational heterogeneity that can elucidate allosteric pathways (9), help explain enzyme catalysis (26), correlate with solution NMR data (27), and reveal new allosteric effects from ligands (16), among other advantages and applications (28–33). Second, to complement these experiments in crystals, we use local hydrogen-deuterium exchange mass spectrometry (HDX-MS), which provides a high-resolution view of backbone amide dynamics and solvent accessibility (34), allowing us to map distinct, mutation-dependent allosteric effects in solution (35, 36). Exploiting the wealth of available ligand-bound structures of PTP1B from crystallographic small-molecule fragment screening experiments (9, 16, 17), we also validate the ligandability of promising allosteric pockets at the most functionally impactful human variant sites.

Overall, this study establishes a unique pipeline from disease-related human genetics to atomic-level allosteric mechanisms for a highly validated drug target enzyme, and lays the groundwork for future drug development at promising new allosteric sites in its 3D structure.

## Results

### Human variants alter phosphorylation levels of PTP1B substrate proteins

We set out to test whether naturally occurring human variants in PTP1B could be used as tools to identify critical residues and regions of the protein that may be more effectively targeted to develop AOM. To prioritize human PTP1B variants for functional characterization, we studied people with the extreme phenotype of persistent healthy thinness (Body Mass Index, BMI <19 kg/m^2^) recruited into the Study into Lean and Thin Subjects (STILTS cohort; www.stilts.org.uk) (37).

Analysis of whole-exome sequencing data on 997 people of UK descent recruited to the STILTS cohort (**Fig. 1A**, **Table S1**) identified 29 unrelated people who carried one of six missense variants (allele frequency, AF<1%) in the gene encoding PTP1B (*PTPN1*) of which two people carried a variant which is rare (AF<0.01%) in population cohorts (Q78R, P302Q; **Table S2**). Among 200,000 unrelated White British exomes from UK Biobank, we identified three predicted protein-truncating PTP1B variants. We also explored whether carriers of any rare missense variants exhibited a trend towards lower or higher mean BMI or proportional BMI categories (BMI >40, BMI >30, BMI <20 kg/m^2^) compared to non-carriers (**Methods**). We selected three additional PTP1B missense variants for exploratory functional characterization: D245G (n=4 carriers; BMI, β (95% confidence interval [CI]) = -5.38 [-9.97, -0.78], glm), L425V (n=4 carriers; β [95% CI] = -4.81 [-9.40,-0.22], glm), and V375M (n=3 of 57 carriers had BMI >40 kg/m^2^; Odds Ratio [95% CI] = 3.1 [0.61-9.4)], Fisher’s exact). Therefore, a total of 12 variants in *PTPN1* encoding PTP1B were taken forward for functional characterization (**Table S4**). Of these, five are located in the ordered catalytic domain, two are in the disordered Pro-rich region, and five are in the remainder of the disordered C-terminal tail or ER anchor (**Fig. 1A**, **Table S2**).

**Figure 1:**
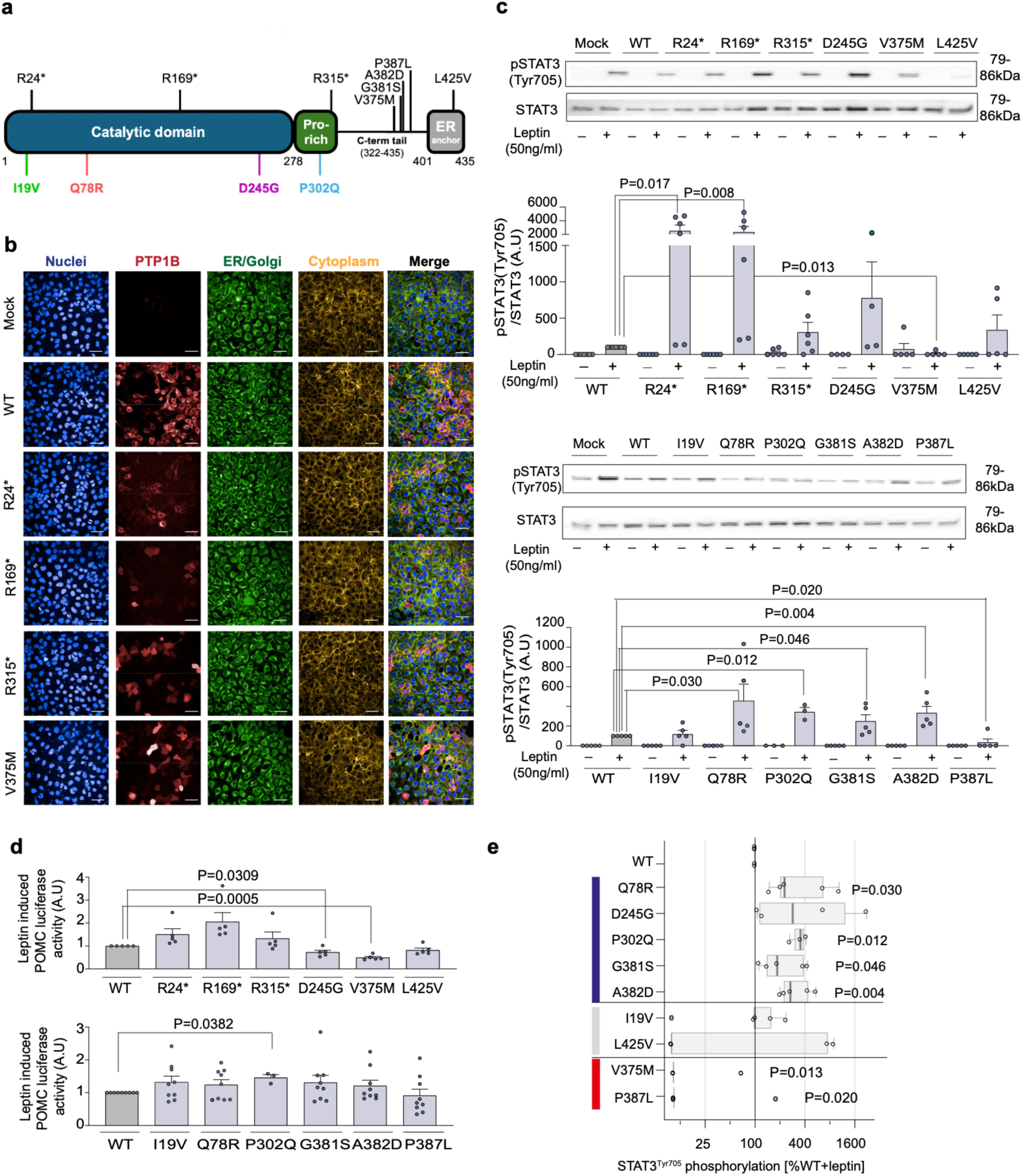
Functional characterization of PTP1B mutants in cells. (A) Mapping of variants identified in the STILTS cohort and UK Biobank on the primary structure of PTP1B (proline (Pro)-rich region; endoplasmic reticulum (ER) anchor). (B) Representative confocal fluorescence microscopy images showing protein localization of WT/mutant PTP1B in HEK293 cells. Blue: DAPI (nuclei), red: Alexa 647 for HA tagged PTP1B, green: Alexa 488 for PDI, yellow: DyLight Phalloidin 554. Scale bar: 50 μm. (C) Effect of WT/mutant PTP1B on leptin-stimulated STAT3 phosphorylation (Tyr705). n=4-7; data expressed as mean +/- SEM normalized to WT (0%) and WT leptin-stimulated (100%) (A.U: arbitrary units). Two-tailed unpaired one-sample t-test on log-transformed data for mutant versus WT. (D) Effect of WT/mutant PTP1B on leptin induced POMC transcription in a luciferase reporter assay. n= 5-9; data expressed as mean +/- SEM relative to WT. Two-tailed unpaired one-sample t-test for mutant versus WT. (E) PTP1B mutations categorized as loss-of-function (LOF; blue), wild-type-like (WT; gray) or gain-of-function (GOF; red) based on phosphorylation and localization assays presented in Fig. 1 and Fig. S1. Statistically significant difference between mutant and WT (expressed as % WT) in leptin-stimulated STAT3 phosphorylation shown. Data are log-transformed; values <10% are set to 10% for visualization. Median shown (box shows interquartile range (IQR); whiskers extend to 1.5*IQR). Results analyzed with an unpaired single-sample t-test (**Table S3**).

To test whether *PTPN1* variants affect the function of PTP1B protein *in vitro*, HEK293 cells were transiently transfected with constructs encoding wild-type (WT) or mutant PTP1B. WT PTP1B was localized to the endoplasmic reticulum (ER) (**Fig. 1B**) and suppressed leptin-dependent phosphorylation of STAT3 and transcription of POMC (**Fig. S1A**). WT PTP1B also decreased basal and BDNF-stimulated phosphorylation of TRKB and insulin-stimulated AKT phosphorylation (**Fig. S1A**).

We investigated whether human PTP1B mutants alter protein expression and/or cellular localization using confocal microscopy of permeabilized transfected cells (**Fig. 1B**, **Fig. S1B-C**). Q78R, P302Q, G381S, and A382D significantly increased leptin-stimulated phosphorylation of STAT3; P302Q PTP1B also potentiated POMC transcription, causing a significant loss of function (**Fig. 1C-D**). Some of these mutants also increased BDNF-stimulated TRKB phosphorylation and enhanced insulin-mediated AKT phosphorylation (**Fig. S1D-E**). In total, 7 of 12 variants studied caused a statistically significant loss of function (LOF) in one or more cellular assays (**Table S4**).

In addition to these LOF variants, V375M PTP1B was mislocalized to the cytoplasm, decreased leptin-stimulated STAT3 phosphorylation to 86% that of WT PTP1B, and decreased POMC transcription, consistent with a significant gain of function (GOF) (**Fig. 1B-D**, **Fig. S1B**). P387L PTP1B also caused a decrease in STAT3 phosphorylation compared to WT PTP1B (**Fig. 1C,E**), consistent with a GOF, although it did not affect POMC transcription. Thus 2 of 12 variants studied caused a statistically significant gain of function (GOF) (**Table S4**).

### Catalytic domain variants allosterically reduce enzyme activity

Building off these observations of altered PTP1B activity in cells, we next aimed to focus on the variants that were most likely to drive discovery of structurally based allostery and associated downstream drug design. To do so, we interrogated the four missense variants within and near the catalytic domain (**Fig. 2A**) in functional and structural detail. Some of these variants are located near known allosteric sites: e.g. D245G and Q78R are adjacent to the conformationally bistable Loop 16, which is known to be allosterically linked to the bistable, dynamic, catalytically essential WPD loop in the active site (9). The locations of others, such as I19V in the α2’ helix and P302Q in the Pro-rich region immediately C-terminal to the allosteric α7 helix, do not correlate with previously known allosteric mechanisms for PTP1B, indicating that further biophysical investigations would be needed.

**Figure 2:**
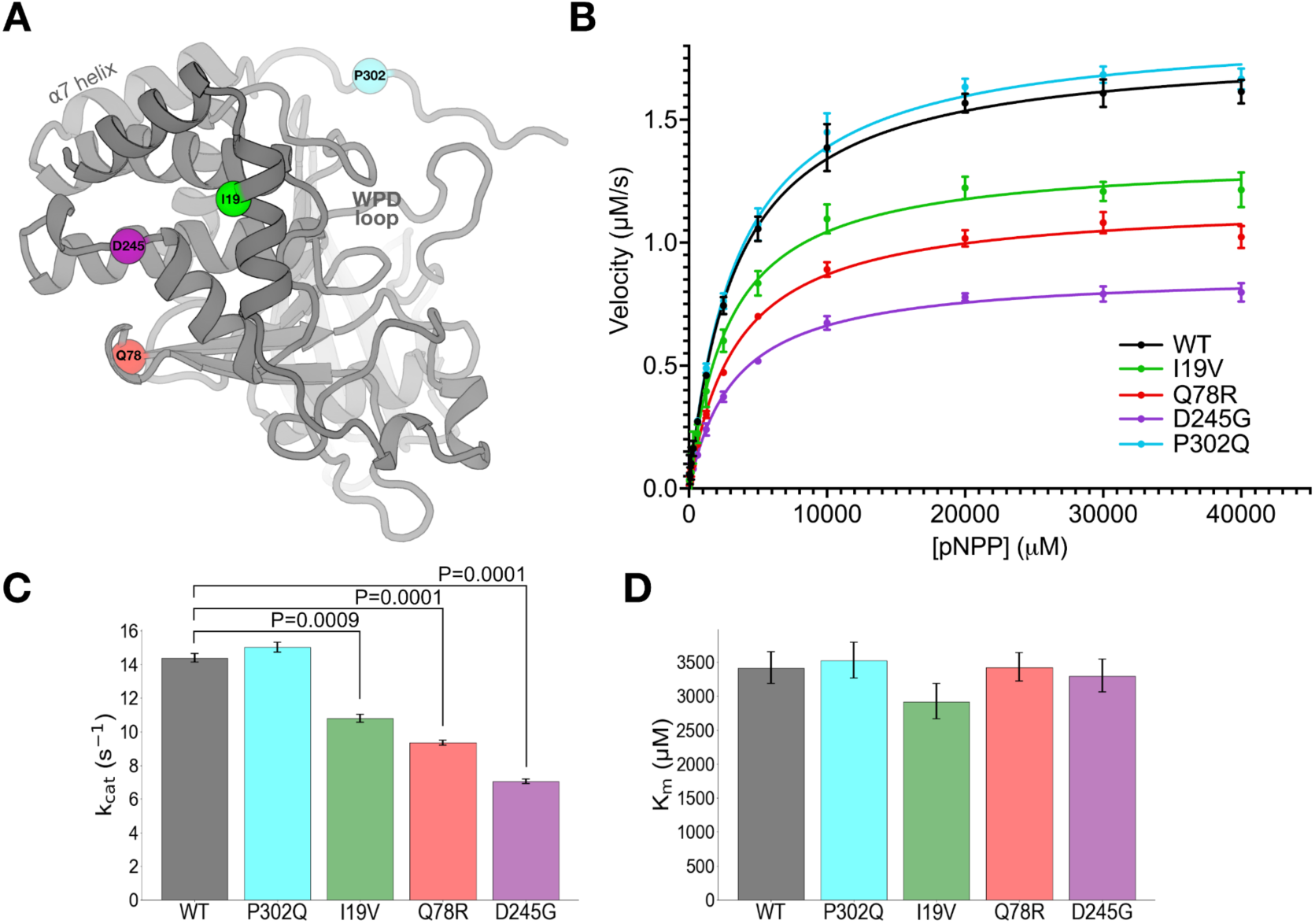
Mutations distal from the active site allosterically perturb enzyme activity *in vitro*. (A) Locations of clinically observed mutations in/near the PTP1B catalytic domain mapped to an AlphaFold 2 (38) structural model obtained from the AlphaFold Database (39). Active-site WPD loop and allosteric α7 helix are labeled. (B) Michaelis-Menten enzyme activity assays using *para*-nitrophenyl phosphate (pNPP) for mutants vs. WT. Error bars represent 95% confidence intervals. (**C**-**D**) Kinetics parameters from Michaelis-Menten analysis. Error bars represent the bounds of the 95% confidence interval based on 8 replicate results. Statistical significance of mutant vs. WT differences was obtained from two-tailed unpaired one-sample t-tests. (C) k_cat_ decreases for several mutants, but (**D**) K_m_ is unchanged (all differences between values are not statistically significant at P=0.05).

To study these four variants in more detail, we first performed *in vitro* enzyme activity assays with a purified recombinant enzyme construct containing the catalytic domain plus disordered Pro-rich region, which is the most commonly employed construct for PTP1B biochemistry and structural biology (residues 1–321; see Methods) (**Fig. 2B**). Compared to WT, D245G and Q78R significantly decreased catalysis *in vitro*, consistent with them having the most extensive LOF effects in cells (**Table S4**). Somewhat surprisingly, I19V also decreased catalysis *in vitro*, despite having no statistically significant effects in cells. These mutations decrease k_cat_ but do not change K_m_ (**Fig. 2C-D**), consistent with allosteric effects from their locations distal from the active site (**Fig. 2A**). In contrast to the other catalytic domain mutations, P302Q had no effect on catalysis *in vitro* (**Fig. 2B-D**), suggesting that its LOF in cells (**Fig. 1C-E**) may occur by other mechanisms such as altered protein-protein interactions in the cellular environment.

To complement the activity assays, we predicted the impact on protein stability for each variant. Mutational analysis with FoldX (40) using several different input structures in different conformational states suggests that D245G is consistently destabilizing (ΔΔG > 3.2 kcal/mol) (**Table S5**). Thus D245G may cause local unfolding that biases the equilibrium of Loop 16 and allosterically modulates the distal active-site WPD loop. By contrast, the other mutations in/near the catalytic domain have less impact on predicted overall stability (**Table S5**), and may affect activity by other, subtler allosteric means.

### Room-temperature crystallography reveals conformational heterogeneity linking variant sites to the active site

To attribute more detailed structural mechanism to these catalytic effects, we used room-temperature (RT) X-ray crystallography. Compared to traditional cryogenic-temperature crystallography, RT crystallography reveals elevated protein conformational heterogeneity (41) which has been shown to underlie biological functions including enzyme catalysis and allosteric signaling (9, 16, 26, 42–46).

The mutations were evident in 2Fo-Fc and Fo-Fc electron density maps for D245G and I19V (**Fig. S2**). Difference density for the Q78R mutation was less clear, likely due to this residue’s high surface accessibility and this structure’s relatively lower resolution (**Table S6**). For D245G in particular, the difference density reveals correlated disappearance of the D245 side chain, appearance of a new ordered water molecule in its place, and perturbation of the neighboring K247 side chain (**Fig. 3D**).

**Figure 3:**
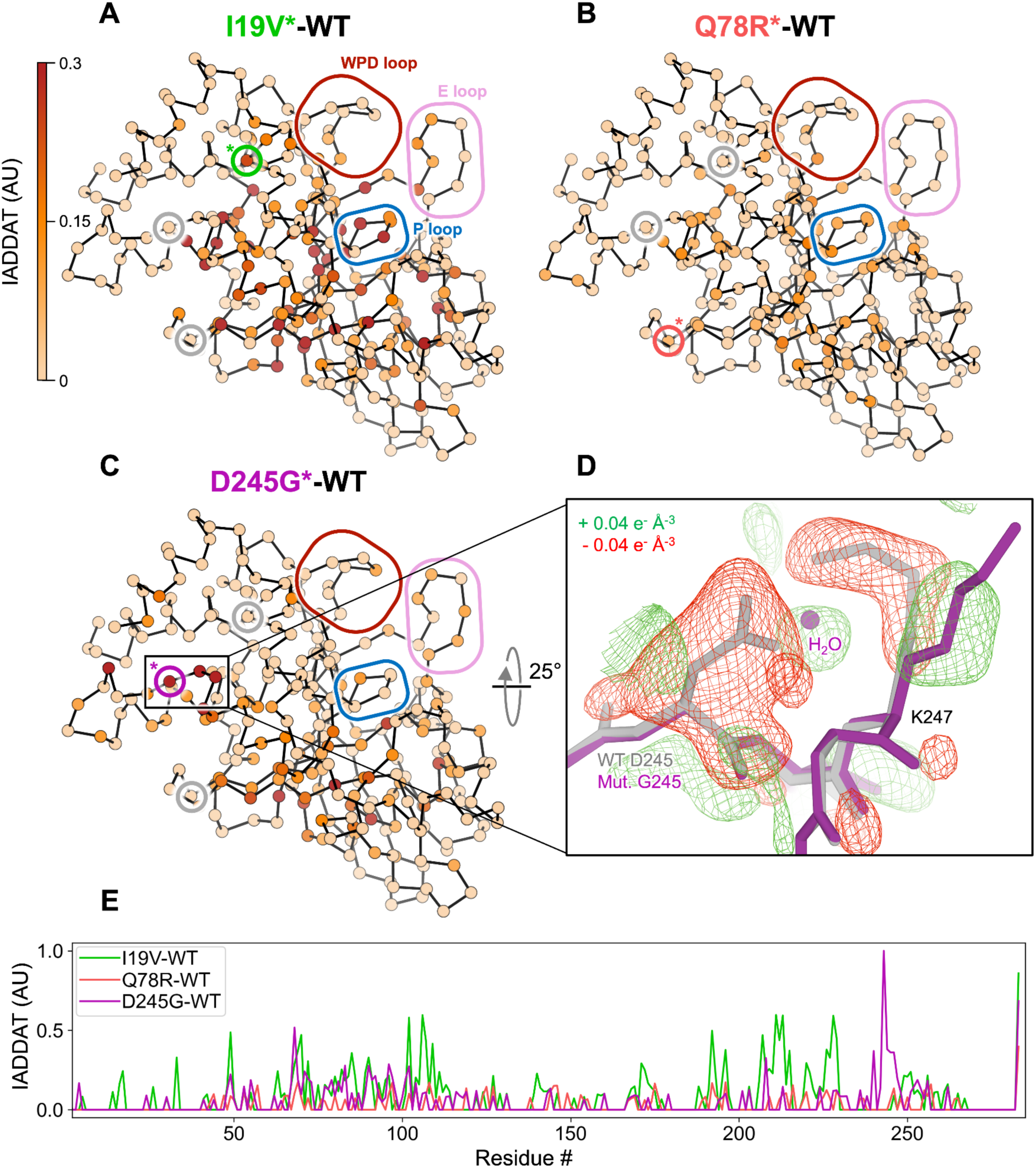
Variants throughout the catalytic domain alter the conformational ensemble of PTP1B in crystals. (**A**-**C**) Weighted isomorphous difference electron density map (*F*_mut_ - *F*_WT_) features were spatially characterized by integrating the absolute difference density above a noise threshold (IADDAT) (47) of 0.04 e^-^/Å^3^ for each pair of mutant vs. wild-type PTP1B room-temperature (RT) crystal structures. IADDAT values were averaged per residue and mapped onto the refined RT mutant structures (Cα positions shown as spheres). IADDAT values range from 0 to 1 (arbitrary units, AU) and are capped at 0.3 for visualization. Small colored circles with a colored asterisk highlight the Cα location for each respective mutation (I19V, green; Q78R, red; D245G, purple; circles for other mutations in gray). Large colored rounded boxes highlight key regions of PTP1B. (A) I19V shows strong IADDAT for the mutation site (marked with an asterisk inside the inlay box), parts of the central β sheet, and active-site P loop. (B) Q78R shows more limited IADDAT, likely due to the lower resolution. (C) D245G shows strong IADDAT for the mutation site, residues in the α4 helix, parts of the central β sheet, and residues near the active site. (D) Weighted D245G-WT difference electron density (+/- 0.04 e^−^ Å^−3^, green/red), focused on the mutation site, overlaid with wild-type (gray) and D245G mutant (purple) structural models. Surrounding protein and solvent atoms that respond to the mutation are highlighted. (E) Plot of per-residue IADDAT vs. sequence for I19V-WT (green), Q78R-WT (red), and D245G-WT (purple).

To quantitatively map detailed, longer-range effects of the mutations on the conformational ensemble of PTP1B, we used integration of absolute difference density above threshold (IADDAT) analysis, as developed previously for time-resolved crystallography (47). With this approach, we observed evidence for conformational disturbances upon mutation that are widespread in the PTP1B structure (**Fig. 3**). For I19V, IADDAT difference features span the mutation site, central β sheet, and active-site P loop (45) (**Fig. 3A**). For D245G, difference features exclude the I19 area, but otherwise span a similar region of PTP1B, including the D245G mutation site and parts of the β sheet and active site (**Fig. 3C**). For Q78R, difference features include some similar regions such as the β sheet and P loop but are more limited (**Fig. 3B**), perhaps due to the lower resolution (**Table S6**). Despite some similarities between these difference maps for these three mutants, in each case the difference density is minimal at the other mutation sites (gray circles in **Fig. 3A-C**), suggesting that each mutation affects the conformational landscape of PTP1B via a distinct allosteric tendril.

### Hydrogen-deuterium exchange maps variant-driven allosteric signals

RT crystallography suggests the mutations have structurally distributed effects on PTP1B in the crystal lattice, but is restricted to the crystalline environment. To assess whether the mutations also affect the structural dynamics of PTP1B in solution, we used high-resolution local hydrogen-deuterium exchange mass spectrometry (HDX-MS) (36). Local HDX-MS measures the relative exchange of labile backbone amide hydrogens at many overlapping peptide sites in a protein, as a proxy for conformational dynamics (34). Using HDX-MS, we obtained peptide maps of exchange with high (∼98.8%) shared coverage of the 1–321 PTP1B sequence across multiple time points for WT and all four mutants in/near the catalytic domain, allowing us to calculate detailed mutant-WT difference Woods plots (**Fig. S3**-**6**).

As visualized by peptide “strip” plots, the four mutations have distinct effects on local conformational dynamics throughout the PTP1B catalytic domain (**Fig. 4A**). Mapping HDX difference values to the protein structure provides further insights into apparent allosteric pathways that may enable the functional effects of the mutations (**Fig. 4B-D**). For example, I19V decreases exchange in the N-terminal α1’ helical region and many other peptides throughout the protein but has little effect on exchange at the active site (**Fig. 4B**), consistent with its weaker catalytic effect *in vitro* (**Fig. 2**) and insignificant functional effects in cells (**Fig. 1**). By contrast, Q78R decreases exchange for an adjacent buried β strand and the active-site P loop, which houses the strictly conserved catalytic cysteine (C215) (**Fig. 4C**). These regions thus form a conduit from the mutation site to the catalytic center, consistent with significant functional effects for Q78R *in vitro* (**Fig. 2**) and in cells (**Fig. 1**). Additionally, D245G increases exchange markedly for the mutation locus itself (**Fig. 4D**), consistent with FoldX computational predictions and RT crystallography (**Fig. 3C-D**). It also decreases exchange dramatically for the active-site pTyr recognition loop (**Fig. 4D**), consistent with D245G exhibiting the most extensive functional impacts *in vitro* (**Fig. 2**) and in cells (**Fig. 1**).

**Figure 4:**
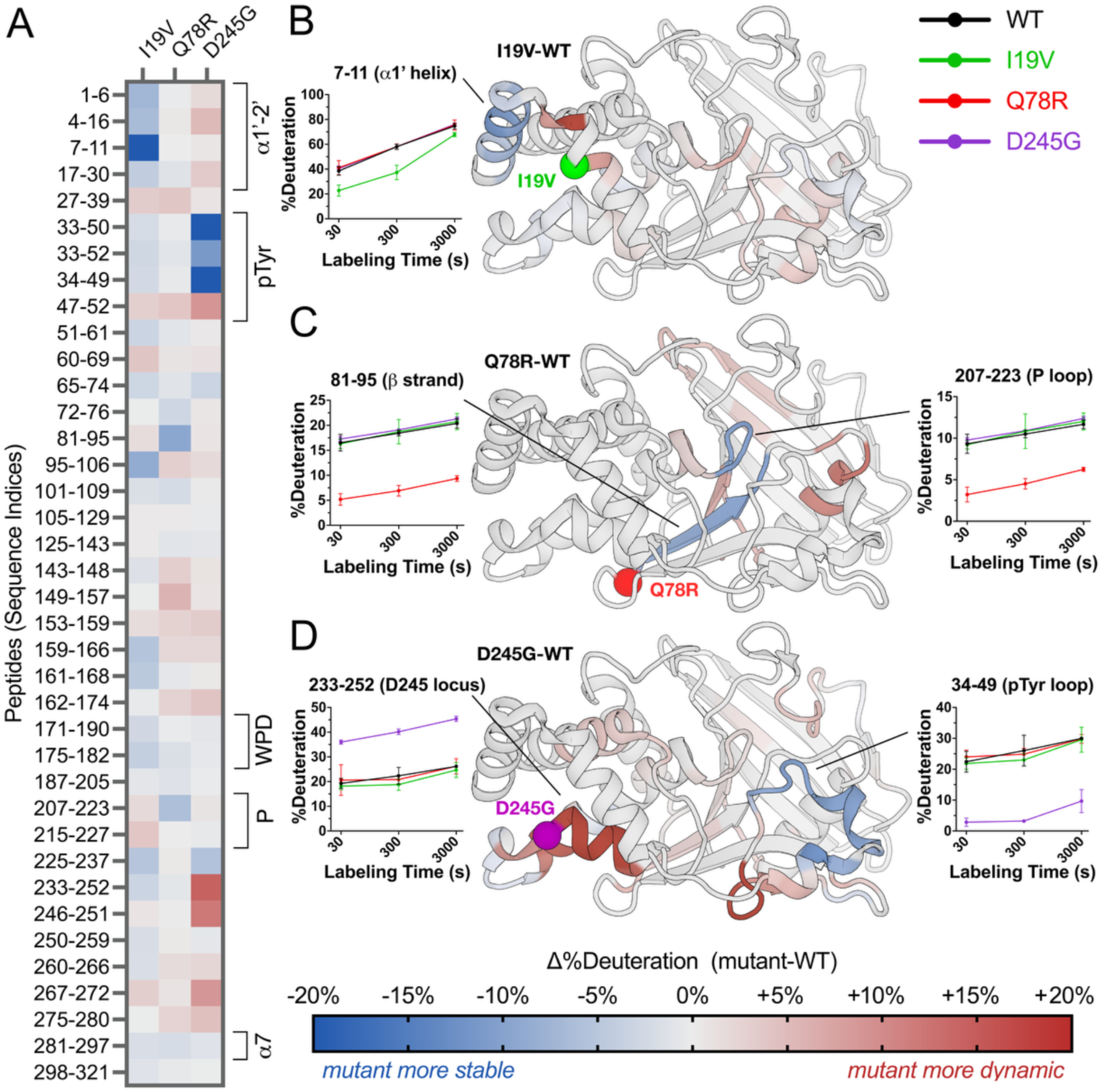
Mutations have widespread effects on protein dynamics in solution by HDX-MS. (A) Mutant-WT difference HDX values at 300 seconds of labeling for selected high-quality peptides spanning the PTP1B sequence. Several key structural regions of PTP1B are indicated with brackets. See color bar for corresponding mutant-WT difference HDX values. (**B**-**D**) Mutant-WT difference HDX values at 300 seconds of labeling at the single-amide level derived from all peptides (see Methods) mapped to a crystal structure of WT PTP1B (PDB ID: 1T49 (25)) for (**B**) I19V, (**C**) Q78R, and (**D**) D245G. For P302Q, see **Fig. S7A**. Residues with Δ%deuteration values between -5% and +5% are colored gray for visual clarity. Deuteration buildup over the time course is shown for select peptides.

### A combined allosteric network from crystallography and HDX-MS

RT crystallography and HDX-MS monitor fundamentally different physicochemical phenomena: crystallographic conformational heterogeneity captures both static and dynamic disorder across a broad range of timescales from picoseconds to seconds or longer; by contrast, hydrogen-deuterium amide exchange measures local backbone solvent accessibility and hydrogen bonding dynamics, primarily detecting structural fluctuations and local unfolding events on the millisecond-to-second timescale. To assess the degree to which these different experiments capture different aspects of the allosteric wiring within the PTP1B fold, we pooled residues affected by any of the three catalytic domain mutations as seen by RT crystallography (**Fig. 5A**) or HDX-MS (**Fig. 5B**). The structural distributions are noticeably different: residues with high RT IADDAT cluster in the center of the protein, whereas residues with high difference HDX values (from selected peptides; see Methods) form two distinct clusters: one in the α-helical bundle (left in **Fig. 5B**) and another in the pTyr recognition loop (right in **Fig. 5B**).

**Figure 5:**
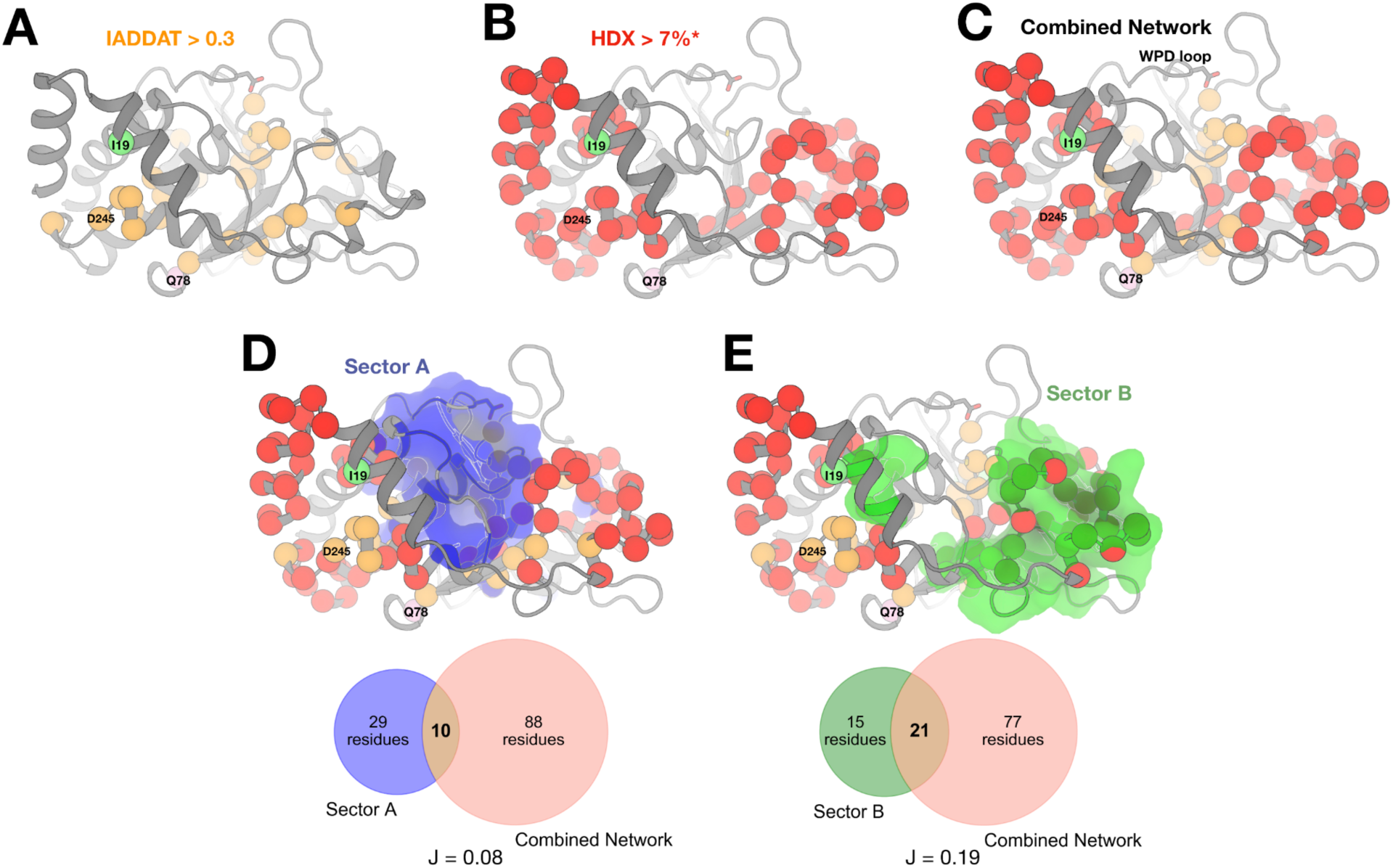
Comparison of networks from RT crystallography, HDX-MS, and coevolving sectors. Each panel maps the pooled set of residues with high response from RT crystallography and/or HDX-MS for all three catalytic domain variants (I19V, Q78R, D245G) to a representative structure of the catalytic domain of PTP1B (PDB ID: 1SUG). The variant sites and the catalytic WPD loop are labeled. (**A**) Residues with RT crystallography IADDAT values > 0.3 (arbitrary units; see **Fig. 3**) are shown as orange spheres. (**B**) Residues with HDX-MS absolute Δ%deuteration > 7% (see **Fig. 4**) are shown as red spheres. * = selected peptides chosen to maximize total sequence coverage with minimal peptide sizes (see Methods). (**C**) The combined network of residues with (A) high IADDAT and/or and (B) high absolute Δ%deuteration. (**D-E**) Coevolving sectors A (blue) and B (green) (24) compared to the combined network. Venn diagrams show the overlap between the sectors and the combined network, with each Jaccard ratio (ratio of intersection to union; J(X,Y) = |X⋂Y| / |X∼Y|) indicated below. Hypergeometric tests were performed to assess overlap between the combined network and each sector, resulting in P = 0.98 for sector A (not statistically significant) and P = 0.011 for sector B (statistically significant, P < 0.05).

When the sets of residues affected in both experiments are pooled, the combined network spans the two HDX clusters via the IADDAT cluster, forming a more cohesive, interconnected network (**Fig. 5C**). The central bridge region spans the β1-β2 hairpin that includes Q78 (**Fig. 5C**). This bridge involves IADDAT contributions from I19V and D245G in the β1 strand as well as HDX contributions from Q78R in the β2 strand. Previously for apo WT PTP1B, β1 was reported to exhibit higher-than-expected amide exchange given its presence in secondary structure and its low variability among crystal structures, possibly due to its location as a solvent-exposed edge strand (36), consistent with its flexibility observed here in response to mutations. Thus, these two techniques appear to offer complementary views of the inherent allosteric network in PTP1B that is sensitive to perturbations by mutations at multiple sites.

We next sought to determine whether the allosteric network identified in our study corresponds to previously characterized allosteric networks in PTP1B or PTPs more broadly. Prior statistical coupling analysis of multiple sequence alignments identified two predominant sectors of coevolving residues in PTPs: sector A, which encompasses the WPD loop and other catalytic elements, and sector B, whose functional role remained poorly understood (24).

Sector A residues, including the WPD loop, have been shown to participate in allosteric mechanisms whereby ligand binding results in large-scale conformational shifts between open and closed states, exemplified by BB3 locking the loop in an open conformation. Notably, our combined network, derived from multiple mutational and structural biology experiments, shows minimal overlap with Sector A (**Fig. 5D**). This suggests that the allosteric mechanisms engaged by the human variants studied here do not conform to the established WPD-loop-mediated pathways of sector A.

Instead, our network exhibits approximately twofold greater, statistically significant overlap with sector B, despite sector B being slightly smaller than sector A (**Fig. 5E**). This overlap is particularly pronounced in the pTyr recognition loop region, as revealed by HDX-MS (**Fig. 5B**). These findings provide direct evidence that perturbations within sector B can modulate enzymatic activity, establishing it as an active conduit for allosteric regulation rather than merely a structurally dynamic region. Taken together, our results show that the variants in this study engage an allosteric pathway distinct from the WPD-mediated network of sector A, revealing a new mode of activity modulation within the PTP1B fold.

### Allosteric effects in the catalytic domain from a mutation in the Pro-rich region

In contrast to the mutations within the ordered catalytic domain, P302Q in the Pro-rich region adjacent to the catalytic domain had no effect on enzymatic activity *in vitro* (**Fig. 2A**) yet caused a LOF in cells (**Fig. 1C-E**). P302Q is located just beyond the α7 helix (residues ∼284–298) (**Fig. 2B**), which alternates between ordered and disordered states (9) and has a critical allosteric role in phosphatase function for PTP1B (10, 19) and its close homolog TCPTP (48). Immediately C-terminal to α7, residues 300 and beyond are thought to be intrinsically disordered: for example, P302 is unmodeled in all available crystal structures of PTP1B. However, the AlphaFold 2 (AF2) structural model for full-length PTP1B (38, 39) includes a relatively “confident” prediction that residues 300–303 adopt an ordered conformation near the active-site WPD loop (**Fig. 2B**). This conformation conflicts with our crystal form (**Fig. S7B**), indicating that crystallography is not a suitable technique to study P302Q.

We therefore used local HDX-MS to examine the effects of P302Q. Many regions positioned near P302 in the AF2 model undergo increased exchange upon P302Q mutation, most prominently the active-site E loop (**Fig. S7A**). By contrast, exchange of other key active-site loops such as the WPD loop, P loop, and Q loop are unaffected, consistent with the lack of catalytic effect for P302Q with purified recombinant protein. The regions affected by P302Q are largely distinct from those affected by the other three variants which are located on the other side of the catalytic domain (**Fig. 5**). Together, these results support the AF2 computational model of PTP1B (39). They further suggest that P302Q modulates the PTP1B conformational ensemble in ways that affect function in cells but do not affect inherent catalysis (such as altering recruitment of various polypeptide substrates) via structural mechanisms that may involve the E loop, which is known to undergo correlated motions with the catalytic WPD loop in another PTP (49).

### Allosteric ligand hotspots colocalize with top human variants

The above analyses focused on characterizing the cellular, biochemical, biophysical, and structural effects of human variants in the enzyme PTP1B. Genetic variants can be used as a tool to inform drug discovery by indicating allosteric weak points on a protein that can be targeted in the drug design process. With this view in mind, we explored hundreds of crystal structures of PTP1B with different small-molecule ligands from the Protein Data Bank (PDB) (17), including high-throughput fragment screens (9, 16), to see if ligand binding coincides with our variant sites.

Indeed, the pockets near I19V, Q78R, and D245G are bindable by dozens of small-molecule fragments, as well as small-molecule buffer components that became fortuitously ordered in crystals (50) (**Fig. 6**). Based on specialized electron density maps that reveal low-occupancy binding events and protein responses in atomic detail (51), we identified one of these fragments that allosterically shifts the conformation of the catalytic WPD loop from open to closed (17).

**Figure 6:**
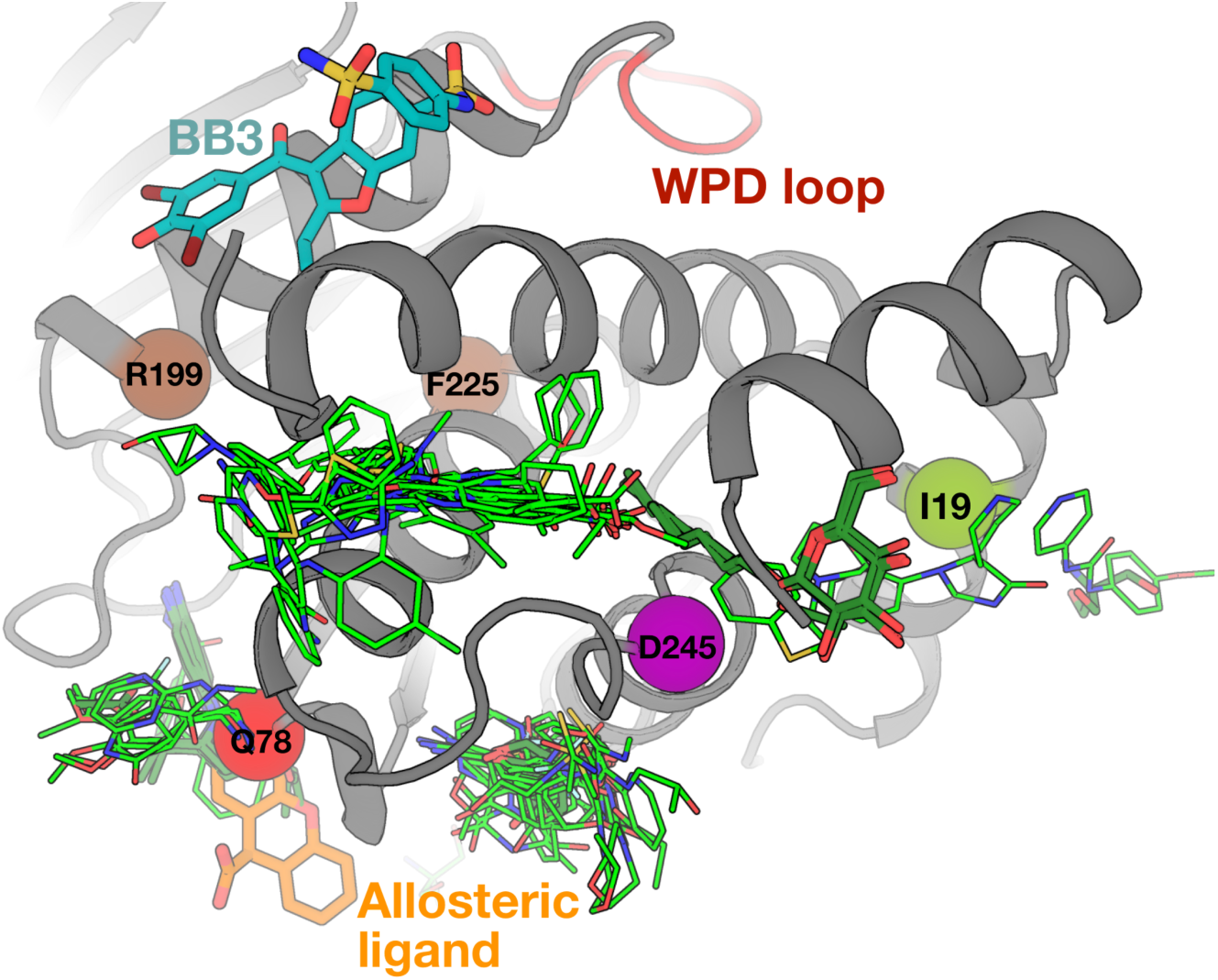
Allosteric sites revealed by human variants are highly ligandable. Ligand binding seen in crystallographic small-molecule fragment screens (light green) (9, 16) and from other structures of PTP1B in the PDB (dark green) indicate significant ligandability at sites near the mutations highlighted in this paper. This includes one fragment (orange, PDB ID: 7GTV) in a site that was shown to allosterically modulate the active-site WPD loop conformation in specialized electron density maps (top, red) (17). The mechanistically distinct BB allosteric site (cyan) (25) and two residues at which mutations were shown to allosterically activate PTP1B activity (brown) (11) are shown for context.

Notably, its binding site is adjacent to one of the mutations with significant functional impacts, Q78R (**Fig. 6**). While this fragment does not inhibit WT or mutant PTP1B with *in vitro* assays (data not shown), this is unsurprising because small-molecule fragments typically bind only weakly and require further optimization for functional modulation. Taken together, the extensive ligand binding coverage near these three functionally impactful mutations in PTP1B suggests that human amino acid variants can be leveraged to pinpoint potentially druggable allosteric sites.

## Discussion

Here we have studied the human PTP1B protein, which, based on a wealth of preclinical evidence regarding its mechanism of action, is a target for weight loss therapy. Our experiments in cells establish that several human variants in *PTP1B* cause a significant LOF and two cause a GOF, illuminating how PTP1B structural elements and domains beyond the active site regulate cellular localization and enzymatic function (**Table S4**). Focusing on allosteric effects in the ordered catalytic domain, we show that different variants perturb the enzyme’s conformational ensemble in distinct but overlapping ways. Moreover, several of the most impactful variants colocalize with ligand-binding hotspots. Together, these molecular and structural findings underscore the promise of targeting specific allosteric sites distal to the central catalytic machinery with small-molecule inhibitors for PTP1B (9, 18, 20, 25).

Although we primarily focused on catalytic domain mutants, several of the PTP1B variants are located in the intrinsically disordered proline-rich region and C-terminal tail (**Fig. 1A**). These regions of PTP1B are absent from crystal structures but can modulate phosphatase function through through a variety of mechanisms (52, 53) including binding to SH3 domains (54), intracellular localization to the ER (55), calpain proteolysis (56), and serine phosphorylation (54), which may be disrupted by the mutations. P302Q lies within the LxVP motif (299-LEPP-302). It may disrupt binding with the Grb2 adaptor protein (57), as suggested by HDX-MS data (**Fig. S7A**) showing increased exchange in the predicted proximal regions of the catalytic domain, consistent with local structural disruption. P387L lies within the SPxK motif (383-QAASPAK-389), implicated in Cdc14 and CDK-mediated S386 phosphoregulation, suggesting that this mutation interferes with phosphorylation-dependent regulation. Although the C-terminal domain is mostly disordered, specific residues such as V375 exhibit partial helical character, as observed by NMR experiments (26). V375, R371, and R373 undergo significant chemical shift perturbations upon binding the allosteric inhibitor MSI-1436, suggesting allosteric communication (19). Although detailed structural ensembles of the disordered C-terminal region are challenging to characterize, proteomics studies comparing these variants to WT PTP1B (57) could clarify how they alter substrate recruitment and specificity, offering insights into endogenous regulatory mechanisms that govern PTP1B activity.

RT crystallographic IADDAT and solution HDX-MS provide complementary perspectives on how human variants perturb the structural dynamics and catalytic efficiency of PTP1B. For each human variant reported here, different structural regions are implicated by IADDAT vs. by HDX-MS (**Fig. 3**, **4**). Combining results from both types of experiments yields a more cohesive, interconnected allosteric network (**Fig. 5**). These observations are in line with previous work showing both similarities and differences between (i) flexibility among a pseudo-ensemble of many crystal structures in different experimental conditions (58–60) and (ii) HDX in solution (36). Moving forward, integrative modeling approaches that incorporate multiple complementary experimental and computational techniques, such as HDX-MS and molecular dynamics simulations for protein-solvent interactions (35), aided by emerging HDX-MS data processing algorithms to enable greater spatial resolution and thermodynamic information (61), will be critical for elucidating the multifaceted nature of allosteric communication in proteins.

The complementary structural biology experiments reported here highlight the ability of several variant sites in PTP1B to allosterically affect distal regions including the active site. Importantly, these sites also reside at the protein surface, and direct orthogonal evidence points to their ability to bind many small-molecule ligands (**Fig. 5**). Thus these variant sites meet both key criteria for potentially druggable allosteric sites. Most of the ligands that bind nearby are small-molecule fragments, which bind only weakly (often with K_d_ values in the mM range), yet can be successfully developed into high-affinity binders and potent allosteric modulators by structure-based fragment assembly, merging, and growth (62, 63). In addition to fragments, the potent allosteric inhibitor SHP099 binds to a structurally equivalent site in the PTP1B paralog SHP2 (64) (**Fig. S8**). Although SHP099 operates by a SHP2-specific mechanism involving SH2 regulatory domains absent from PTP1B, it provides complementary evidence of ligandability for this area of the structurally conserved PTP fold (65) that could be exploited for structure-based drug design. Taken together, although designing potent new allosteric inhibitors is beyond the scope of the current report, the allosteric sites uncovered here provide promising new footholds for future development of allosteric inhibitors of PTP1B that operate by distinct allosteric mechanisms from existing compounds.

Effective PTP1B inhibitors could play an important role in weight management. Drugs that improve leptin sensitivity (e.g. withaferin) reduce food intake and body weight in obese mice (characterized by leptin resistance) but not in lean mice (66). Clinical trials will be needed to test whether drugs that inhibit PTP1B and thereby increase the amplitude of leptin signaling have a meaningful impact on body weight, either in people with obesity (which may be characterized by a degree of leptin resistance (67)) or in people in the weight-reduced state, where relative leptin deficiency is a major driver of weight regain (68, 69).

## Materials and Methods

### Ethics

The study was reviewed and approved by the South Cambridgeshire Research Ethics Committee (12/EE/0172). All participants provided written informed consent prior to inclusion.

### STILTS cohort

To recruit people for the Study Into Lean and Thin Subjects (STILTS) cohort we worked in collaboration with 1,143 General Practitioners (GPs) to invite 47,707 eligible people to participate (**Fig. 1A**). Participants of UK European descent, aged 16-65 years and with a BMI <18 kg/m^2^ were invited to take part. We applied strict criteria to exclude participants with medical conditions that can affect body weight, such as chronic renal, liver and gastrointestinal problems, or eating disorders assessed using a validated screening tool (SCOFF) (70). We also excluded participants who stated that they exercised every day, more than 3 times per week or who reported activity levels more than 6 metabolic equivalents (METs) for any duration or frequency. Finally, we only included participants who reported having been thin their whole life. Some participants with a BMI between 18-19 kg/m^2^ were included due to having a strong family history of thinness. One thousand healthy thin participants consented to whole exome sequencing. DNA was extracted from salivary samples obtained using the Oragene 500 kit according to manufacturer’s instructions. We obtained additional data on these individuals using the Three Factor Eating Questionnaire which measures cognitive restraint, uncontrolled eating and emotional eating and has been extensively validated in large cohorts (71). We also obtained data on occupational and social physical activity using a modified version of the EPIC (European Prospective Investigation into Cancer and Nutrition) physical activity questionnaire, which has been validated against direct measurements of energy expenditure including calorimetry and heart rate variability (72).

### Variant detection in the STILTS cohort

Exome sequencing was performed for N=997 people from the STILTS cohort. Batch 1 (N=397 people; BGI Genomics) was aligned to hg19 reference (mean depth of target regions, 71-118X (median, 92X); coverage of target regions >= 20X, 70-94% (median, 91%)) and variants were called using GATK v3.3.0 (BGI Genomics). Batch 2 (N=598 people; Macrogen Inc) was aligned to hg38 reference (mean depth of target regions, 102-167X (median, 148X); coverage (%) of target regions with >= 20X, 94-98% (median, 97%) and variants were called using GATK v4.0.5.1 (Macrogen Inc). Variant filtering was based on GATK Best Practices with hard-thresholded variant filters (SNPs: QD<2, FS>60, MQ<40, MRankSum<-12.5, ReadPosRankSum<-8; indels: QD<2, FS>200, ReadPosRankSum<-20). Variant consequence was obtained from Ensembl VEP (Ensembl v96) with respect to PTP1B (gene *PTPN1*) transcript ENST00000371621 and TCPTP (gene *PTPN2*) transcript ENST00000309660. All PTPN1 (PTP1B) and PTPN2 (TCPTP) missense variants from STILTS exomes were taken forward for molecular functional characterization (**Fig. 1A**, **Table S2**).

### PTP1B variant selection from UK Biobank 200K exomes

This research was conducted using the UK Biobank Resource (project 53821). The pVCF file for chromosome 20, block 17 was obtained from UK Biobank OQFE 200K exomes interim release (Field 23156). Multiallelics were split and left-normalized. Variants were annotated using Ensembl VEP (v96 cache; GRCh38) and filtered to retain variants with IMPACT=HIGH or MODERATE with respect to PTP1B (gene *PTPN1*) transcript ENST00000371621. Samples were filtered to retain European exomes (Field 22006; self-reported “White British” and tight cluster in genotype PCA). Since kinship was not modeled in this analysis, related pairs up to third-degree kinship were obtained from the UK Biobank Genetic Data resource (ukb_rel.dat) and one person was excluded from each related pair (third-degree kinship). UK Biobank phenotypic and genetic data were obtained for BMI (field 21001.0.0), age, sex, genetic principal components, and WES sequencing batch (50K or 150K).

We identified predicted protein-truncating variants (Ensembl VEP Impact=HIGH, which detected three stop-gain mutations) and, with the aim of augmenting the *PTPN1* missense variants taken forward for exploratory molecular functional characterization, we explored whether any rare missense variants exhibited a trend towards lower or higher BMI, or proportional BMI categories, among carriers compared to non-carriers among unrelated White British exomes from UK Biobank 200,000 exomes. Exploratory single-variant association tests were performed for BMI phenotypes: (i) Fisher’s Exact test for dichotomized BMI (BMI >40, BMI >30, BMI <20 kg/m^2^) by variant carrier status, (ii) exploratory case-control regression analysis of dichotomized BMI (plink2 –glm with Firth regression) and continuous BMI (plink2 –glm) with covariates age, sex and forty genetic principal components (Fields 22009.0.1-40), together with inspection of the cumulative distribution of BMI and adjusted BMI among variant carriers and non-carriers. Three *PTPN1* stop-gains and three missense variants from UK Biobank 200K exomes were taken forward for exploratory functional characterization (**Table S2**, **Fig. 1A**).

### Cell culture

HEK293 cells were cultured in high glucose Dulbecco’s modified eagle medium (DMEM, GIBCO, 41965) supplemented with 10% fetal bovine serum (GIBCO, 10270, South America origin), 1% GlutaMAX (100X) (GIBCO, 35050), and 100 units/mL penicillin and 100 μg/mL streptomycin (Sigma-Aldrich, P0781).

### Cloning of PTP1B variants for in-cell assays

PTP1B cDNA constructs containing an N-terminal HA tag in pCDNA3.1 (+) vector and TCPTP cDNA constructs containing a C-terminal DYK tag in pCDNA3.1 (+) vector were used throughout the study. Vector containing the N-terminal HA-tagged wild-type (WT) PTP1B (GenScript OHu27552C) and vector containing the C-terminal DYK-tagged wild-type (WT) (GenScript OHu17864D) were used for site-directed mutagenesis using Q5 site-directed mutagenesis kit (NEB, E0554S) according to the manufacturer’s protocols. All constructs were verified with Sanger sequencing.

### Cell transfection and stimulation

HEK293 cells were seeded in 96-well plates coated with Poly-D-Lysine (20,000 cells/well). For STAT3 phosphorylation experiments cells were transiently transfected with 100 ng/well plasmid encoding either empty pcDNA3.1(+) vector (negative control), WT or mutant PTP1B or TCPTP plasmid, combined with 50 ng/well plasmid for Leptin Receptor, and 10 ng/well plasmid for STAT3 using Lipofectamine 2000 in Opti-MEM I medium according to the manufacturer’s protocols. For TRKB phosphorylation experiments cells were transiently transfected with 100 ng/well plasmid encoding either empty pcDNA3.1(+) vector (negative control), WT or mutant PTP1B plasmid, combined with 25 ng/well plasmid for TRKB Receptor. For AKT phosphorylation experiments cells were transiently transfected with 100 ng/well plasmid encoding either empty pcDNA3.1(+) vector (negative control), WT or mutant PTP1B plasmid using Lipofectamine 2000 in Opti-MEM I medium according to the manufacturer’s protocols. After 5 hours transfection, media was replaced by DMEM. Cells were starved overnight, stimulated with either 50 ng/mL leptin (Millipore, 429700) (STAT3 phosphorylation) for 10 minutes, 50 ng/mL BDNF (R&D, 248-BDB-050/CF) (TRKb phosphorylation) for 5 minutes or 200 ng/mL insulin (Sigma, I9278) (AKT phosphorylation) for 10 minutes.

### Western blotting

After stimulation cells were washed in PBS and lysed in 40 μL radio-immunoprecipitation assay buffer (RIPA) (Sigma, R0278) supplemented with protease and phosphatase inhibitors while on ice. 96-well plates with cells lysed in the RIPA buffer were shaken vigorously for 30 seconds and kept on ice for 1 minute (x3) until all cells were detached. 30 μL per well of RIPA buffer with lysed cells were transferred to 96 well plates combining 2-4 wells per condition. Plates were kept on ice for 10 minutes and then harvested by centrifugation at 4000 rpm for 20 min and prepared for electrophoresis as described by the manufacturer’s protocol using the iBOLT 2 Invitrogen system (B0007, B0009, NW04127BOX, IB23001). Membranes were blocked with 3% BSA solution in TBS-T for 1 hour at room temperature and probed overnight at 4 ^ο^C using Rabbit anti-STAT3 at 1:1000 dilution (Cell Signaling Technology, 12640), Rabbit anti-Phospho STAT3 (pY705) (Cell Signaling Technology, 9145) at 1:1000 dilution, Rabbit anti-Phospho TRKa (pY674/675)/ phospho TRKb (pY706/707) (Cell Signaling Technology, 4621) 1:1000 dilution, Rabbit anti-TRKb (Cell Signaling Technology, 4603) 1:1000 dilution, Rabbit anti-HA (Cell Signaling Technology, C29F4) at 1:1000 dilution, Mouse anti-DYK (FLAG) tag M2 antibody (Sigma, F1804), Rabbit anti-phospho AKT (pS473) (Cell Signaling Technology, 4060) 1:1000 dilution, Rabbit anti-AKT pan (Cell Signaling Technology, 4691) 1:1000 dilution, Rabbit anti-βACTIN (Cell Signaling Technology, 4967) at 1:1000 dilution and Rabbit anti-vinculin (Abcam, ab129002) all prepared in the blocking buffer. Cells were washed three times with TBS-T for 10 min at room temperature with gentle shaking and were incubated with secondary antibody, Goat anti-rabbit IgG-HRP (Dako, P0448) or Goat anti-mouse IgG-HRP (Dako, P0447) diluted 1:2500 in 3% BSA in TBS-T for 1 hour at room temperature. Bands were developed using the SuperSignal West Dura Extended Duration Substrate (ThermoFisher Scientific, 34075) and imaged in either BioRad Chemidoc XRS or Chemidoc MP Imaging (BioRad) according to the manufacturer’s protocols. For total TRKB or AKT, blots were stripped for 15 minutes in 10 mL 1X Re-blot Plus Strong Solution (EMD Millipore, 2504) and blocked with the antibody. The band intensity of western blots was quantified using FIJI. For data normalization in STAT3 or AKT phosphorylation, unstimulated WT PTP1B or TCPTP readouts were set as baseline (0) and maximum WT PTP1B or TCPTP STAT3 or AKT phosphorylation upon stimulation was set as 100%. For data normalization in TRKB phosphorylation, unstimulated WT PTP1B readout was set as 100%. For data normalization in mock versus WT in STAT3 or AKT phosphorylation unstimulated mock (empty vector) readouts were set as baseline (0%) and maximum mock STAT3 or AKT phosphorylation upon stimulation was set as 100%. For data normalization in mock versus WT in TRKB phosphorylation unstimulated mock (empty vector) readouts was set as 100%. All uncropped blots analyzed are part of Fig. S1.

### Luciferase POMC transcription activation assay

HEK293 cells were seeded into white 96-well microplates (chimney well) (Greiner, 655083) coated with Poly-D-Lysine (Sigma, A-003-E) (20,000 cells/well) and transiently transfected the next day with 50 ng/well plasmid encoding either empty pcDNA3.1(+) vector (negative control), WT or mutant PTP1B plasmid, combined with 12.5 ng/well plasmid for Leptin Receptor and 50ng/well plasmid for POMC luciferase using Lipofectamine 2000 (Thermo Fisher Scientific, 11668019) in serum-free Opti-MEM I medium (GIBCO, 31985) according to the manufacturer’s protocols. After 5 hours transfection cells were incubated overnight with starvation media (DMEM no FBS) with or without the presence of 200 ng/ml Leptin (Human Recombinant *E. coli*, EMD Millipore, 429700). Quantitation of firefly luciferase activity was performed using the Steadylite Plus Reporter Gene Assay System (Perkin Elmer, 6066759) according to the manufacturer’s protocol. For data normalization WT PTP1B POMC luciferase readouts were set to 1 and PTP1B mutant values were normalized relative to WT.

### Subcellular localization of human PTP1B mutants

HEK293 cells were seeded in black clear bottom CellCarrier-96 Ultra Microplates (Perkin Elmer, 6055302) coated with Poly-D-Lysine solution (Sigma, A-003-E) (10,000 cells/well) and transiently transfected the next day with 100 ng/well plasmid encoding either empty pcDNA3.1(+) vector (negative control), WT, mutant PTP1B or TCPTP plasmid using Lipofectamine 2000 (Thermo Fisher Scientific, 11668019) in serum-free Opti-MEM I medium (GIBCO, 31985) according to the manufacturer’s protocols. After 24 hours, cells were fixed with 4% Formaldehyde (Fisher Chemicals, F/150/PB17) in Phosphate-buffered saline (PBS) for 20 minutes at room temperature, permeabilized with 0.2% Triton X-100 (BDH, 306324N) for 30 minutes at room temperature, blocked for 1 hour in 3% Bovine Serum Albumin (BSA) (Sigma, A7906) in TBS-T at room temperature. For PTP1B localization, cells were incubated overnight at 4°C with Rabbit anti-HA (Cell Signaling Technology, C29F4) at 1:100 dilution in 3% BSA in TBS-T, Mouse anti-PDI (Thermo Fisher Scientific, MA3-018) at 1:100 dilution, or only 3% BSA in TBS-T as a negative control. Cells were washed three times with PBS for 5 minutes, incubated with goat anti-mouse secondary antibody Alexa Fluor 488 (Thermo Fisher Scientific, A11029) in 1:200 dilution and donkey anti-rabbit secondary antibody Alexa Fluor 647 (Thermo Fisher Scientific, A31573) in 3% BSA in TBS-T for 1 hour at room temperature and washed twice with PBS for 5 minutes, incubated with DAPI (Invitrogen, D1306) in 1:500 dilution in PBS and DyLight 554 Phalloidin (Cell Signaling Technology, 13054) in 1:200 dilution in PBS for 10 minutes, washed once with PBS for 5 minutes and kept in PBS. For TCPTP localization cells were incubated overnight at 4°C with Mouse anti-DYK (FLAG) (Sigma, F1804) at 1:100 dilution in 3% BSA in TBS-T. Cells were washed three times with PBS for 5 minutes, incubated with goat anti mouse secondary antibody Alexa Fluor 488 (Thermo Fisher Scientific, A11029) in 1:200 dilution in 3% BSA in TBS-T for 1 hour at room temperature and washed twice with PBS for 5 minutes, incubated with DAPI (Invitrogen, D1306) in 1:500 dilution in PBS and DyLight 554 Phalloidin (Cell Signaling Technology, 13054) in 1:200 dilution in PBS for 10 minutes, washed once with PBS for 5 minutes and kept in PBS. Cells were imaged in the Opera Phenix High Content Screening Confocal system (Perkin Elmer).

### Quantification and statistical analysis of cellular models

Results were analyzed using GraphPad Prism 8 (GraphPad Software). The difference between mutant PTP1B with WT was estimated and tested using two-tailed one-sample t-tests on original scale or log-transformed data in comparison to either unstimulated or stimulated samples. Nominal p<0.05 was considered statistically significant. In **Fig. 1C**, t-tests were run using WT-normalized data on the original scale (**Fig. 1C**, **Fig. S1D-E**, **Table S3**). In **Fig. 1E**, t-tests were run using log(WT-normalized data) with a zero-value offset of 0.001 (**Fig. 1E**, **Table S3**). Missense variants were considered to be loss-of-function (LOF) if the mean of log(WT-normalized data) differed from log(100%) with p<0.05 in any of the assays for STAT3, AKT or TRKB phosphorylation upon stimulation (**Fig. 1E**, **Table S3**). Studies in cellular models are from at least 3 independent experiments.

### Cloning, expression, and purification of PTP1B variants for biophysical experiments

All biophysical experiments were conducted using wild-type PTP1B sequence with residues 1-321 (9). The construct is housed in a pET24b vector containing a kanamycin resistance gene. In contrast to some previous related crystallography work with PTP1B, a true WT sequence was used here, with no so-called WT* mutations (C32S/C92V) (9). The starting WT construct included amino acid residues 1-435 of PTP1B, then was shortened to residues 1-321 using site-directed mutagenesis. To generate I19V, Q78R, D245G, and P302Q variants, site-directed mutagenesis was employed starting from this 1-321 construct.

The protein expression and purification process followed a previously documented method (73), with some slight modifications. To initiate expression, we transformed plasmids containing the desired site mutation into competent *E. coli* BL21 (DE3) cells. These cells were allowed to incubate on LB + kanamycin plates at 37°C overnight. Subsequently, 5 mL starter cultures of LB and kanamycin at 1 mM final concentration were inoculated with individual colonies and shaken overnight at 37°C. Larger 1 L cultures of LB and kanamycin at 1 mM final concentration were then inoculated with starter cultures and grown with shaking at 37°C until the optical density at 600 nm reached approximately 0.6–0.8. Induction of the 1 L cultures was carried out by IPTG at a final concentration of 500 μM, and shaking was continued overnight at 18°C. Induced cells were harvested by centrifugation, the resulting cell pellets (“cellets”) were flash frozen, and stored at -80°C in 50 mL conical tubes, for subsequent purification.

Prior to purification, harvested cells were resuspended with lysis buffer containing dissolved Pierce protease inhibitor tablets using a vortexer. Resuspended cells were sonicated (on ice) for 10 mins at an amplitude of 50% using 10 s on/off intervals. Cells were then centrifuged and the supernatant syringed filtered with a 0.22 μm filter and reserved for purification. Initial purification involved cation exchange on an SP FF 16/10 HiPrep column (GE Healthcare Life Sciences). This was carried out using a lysis buffer (100 mM MES pH 6.5, 1 mM EDTA, 1 mM DTT) and a NaCl gradient (0–1 M), resulting in protein elution occurring at approximately 200 mM NaCl. Following this step, size exclusion chromatography was conducted utilizing a S75 size exclusion column (GE Healthcare Life Sciences) in a crystallization buffer (10 mM Tris pH 7.5, 0.2 mM EDTA, 25 mM NaCl, 3 mM DTT). The purity of the sample was evaluated through SDS-PAGE analysis, demonstrating it to be devoid of contaminants and displaying a high level of purity.

### *In vitro* enzyme activity assays

To investigate the kinetic parameters of the mutant proteins, a colorimetric assay utilizing *para*-nitrophenyl phosphate (pNPP) as a substrate was executed. The assay buffer was prepared to a final concentration of 50 mM HEPES (pH 7.0), 1 mM EDTA, 100 mM NaCl, 0.05% Tween-20, and 1 mM β-mercaptoethanol (BME), then 0.22 µm filtered and stored at room temperature.

Subsequently, a series of 12 pNPP concentrations, ranging from 40 mM to 3.9 µM, were prepared by serial dilution in the assay buffer to cover a broad range of substrate concentrations for kinetic analysis. Prior to the assay, two independent measurements of each mutant protein’s concentration were performed five times using a NanoDrop One. Afterwards, the mutant protein samples were diluted to matching concentration (250 nM) in assay buffer, and the average protein concentration for each mutant protein was confirmed again.

For the assay, 50 µL of the diluted protein solution was aliquoted into each well of a Corning 96-well flat-bottom non-binding polystyrene plate. The reaction was initiated by adding 50 µL of the pNPP solution to each well, with thorough mixing achieved through gentle pipetting (final protein concentration of 125 nM). Absorbance at 405 nm was measured at 18-second intervals over a 6-minute period using a SpectraMax i3 plate reader, calibrated prior to use.

Each concentration of pNPP was tested in quadruplicate for each mutant protein. The slopes of absorbance change (mAU per minute) over the 6-minute interval were calculated and used to derive the maximum velocity (V_max_) of each reaction. The catalytic constant (k_cat_) was determined by dividing V_max_ by the average concentration of each mutant protein. These k_cat_ values, obtained from two independent experiments, were combined, and analyzed using GraphPad Prism 9 to plot the kinetic curves and to calculate the Michaelis constant (K_m_), with results presented in **Fig. 2**.

We also tested inhibition by a structurally allosteric small-molecule fragment (orange in **Fig. 5**): 9H–xanthene–9–carboxylic acid (PDB: 7GTV). The assay was performed with fragment concentrations from 1–1000 μM, in the same assay buffer as in the kinetics experiments but containing 2% DMSO. Negative controls used only assay buffer plus 2% DMSO, while the potent active-site inhibitor TCS–401 with concentrations from 0.1–120 μM served as a positive control. Each concentration was tested in four replicates, and absorbance slopes were measured as above. Final data were analyzed in GraphPad Prism 9 to compare initial velocities under each condition and, where feasible, derive IC_50_ values.

### Protein crystallization

Purified WT PTP1B mutants were concentrated to a final concentration of about 40 mg/mL before initiating crystallization experiments. The crystallization well solution used in this study consisted of 0.3 M magnesium acetate, 0.1 M HEPES pH 7.5, 0.1% β-mercaptoethanol, 13.5% PEG 8000, 2% ethanol. Crystallization drops were prepared using EasyXtal 15-Well trays (Nextal, 132006). A 1:1 protein-to-solution ratio was employed such that 1 μL of protein solution was mixed with 1 μL of well solution for one set of drops, and 2 μL of protein solution was mixed with 2 μL of well solution for another set of drops. The prepared crystallization trays were then incubated at a controlled temperature of 4°C. Within a short span of approximately 2 days, crystals became evident in the trays. These initial crystals progressively grew over the course of about 1 week, reaching their maximum size during this period. The fully developed crystals obtained final dimensions ranging from approximately 50 х 50 х 100 μm to 300 х 300 х 1000 μm. I19V crystals were generally the largest, and Q78R crystals generally the smallest.

### X-ray diffraction

Crystals for room-temperature (RT) X-ray data collection were harvested under a humidifier using MiTeGen microloops of relevant size and stored in SSRL *In-Situ* Crystallization Plates, allowing crystals to be shipped in humidity controlled chambers for remote data collection at elevated temperature. X-ray diffraction data were collected remotely at the Stanford Synchrotron Radiation Lightsource (SSRL) beamline 12-1. Single crystals were exposed to X-rays at an elevated cryostream temperature of 298 K, with 0.5° of helical crystal translation per image and 0.1 seconds exposure per image, for a total of 180° across 360 images. The unattenuated X-ray flux was measured as 2.0 x 10^12^ photons/second, and the X-ray beam attenuated to between 1–5% transmission.

### Crystallographic data reduction and refinement

The data reduction pipeline fast_dp (74) was used for initial bulk data reduction in order to determine the highest-resolution datasets to process. Selected datasets were reprocessed using the xia2 DIALS (75) pipeline. Ideal resolution cutoffs were automatically determined based on a CC_1/2_ (76) of ∼0.3 and an outer shell completeness of >97%. Molecular replacement and initial refinement were performed using MOLREP and Refmac5 from the CCP4 suite of crystallographic programs (77), using PDB entry 1SUG as the initial molecular replacement search model. Subsequent iterative rounds of refinement were performed using phenix.refine (78) and Coot (79). Hydrogen atoms were added using phenix.ready_set (80, 81). Final PTP1B models were assessed statistically using MolProbity (82). Data collection and refinement statistics are shown in **Table S6**.

### Density map analysis

In order to reduce the noise in our difference electron density maps, we reweighted our maps using the following method. PTP1B mutant unmerged data from DIALS were run through the data reduction program Aimless (83) in order to rescale the data to a reference dataset. In this case, PTP1B WT experimental data (structure factor amplitudes only) were utilized as reference data to solve indexing ambiguity and space group, and to provide a free reflections set for calculating R_free_. Fo-Fo difference structure factors were then weighted using custom scripts within the reciprocalspaceship suite (84), with map phases calculated using a PTP1B WT model refined against structure factor amplitudes (*F*) only. Structure factors were then converted to difference maps using reciprocalspaceship using weights with an alpha value of α = 0.05 to decrease the influence of measurement errors on structure factor difference amplitudes.

To characterize our difference electron density map features across the structure of our PTP1B mutants, we integrated the absolute difference density above a noise threshold (IADDAT) (47). Here, difference peaks with an absolute value greater than 0.04 e^-^ Å^-3^ (converted from equivalent RMSD values in Coot) and within 1.5 Å of a protein heavy atom (excluding waters) were summed and averaged on a per-residue basis. These values were subsequently mapped back onto the corresponding Cα positions in the protein crystal structure. This was achieved using custom scripts within reciprocalspaceship. Jupyter notebooks for structure factor reweighting and IADDAT calculations are available at GitHub repositories associated with reciprocalspaceship (84) and Wolff et al. (47) respectively.

### HDX-MS data collection

#### Sample handling

HDX-MS data collection and processing were performed as recently reported (36) with some minor variations. For each experiment, liquid handling was performed by the LEAP HDX platform. This robotic system precisely initiates and times labeling reactions (reaction volume of 50 µL), followed by rapid mixing with quench solution and dropping the temperature to 0–4°C. Once the sample was thoroughly mixed with quench, 100 µL of this quenched sample was then injected into the pepsin column.

#### Digestion optimization

Digestion optimization experiments were conducted to determine the optimal quench conditions to halt hydrogen-deuterium exchange and prepare the protein for digestion over a pepsin column by inducing partial or extensive denaturing / unfolding. A series of quenching solutions consisting of 1.5% formic acid, 3.0% acetonitrile, and varying concentrations (0, 0.5, 1.0, 2.0, 4.0 M) of guanidinium hydrochloride (GuHCl) was mixed in a 1:1 ratio with the unlabeled protein. The deuteration buffer was identical to the protein buffer used in the non-deuterated experiments, except for the presence of deuterium in high (>99.5%) abundance. To perform local hydrogen-deuterium exchange (HDX) experiments, a Waters Enzymate BEH Pepsin Column was employed to generate peptides for subsequent analysis. Peptic peptides were eluted through a C18 analytical column (Hypersil Gold, 50 mm length × 1 mm diameter, 1.9 μm particle size, Thermo Fisher Scientific) into a Bruker maXis-II ESI-QqTOF high-resolution mass spectrometer. Peptide maps were generated for PTP1B for each of the varied GuHCl conditions. The best coverage and highest resolution were seen with conditions 2.0 M and 4.0 M. A concentration of 3.0 M of GuHCl was chosen for future experiments.

#### HDX labeling

Purified protein samples (PTP1B 1-321 WT, I19V, Q78R, D245G, P302Q) were prepared as described in the previous section. All experiments were carried out in the crystallization buffer at 15°C to best match pH, salt, and reducing conditions present in comparable structural studies of PTP1B. The protein sample was first diluted to 20 µM in the H_2_O crystallization buffer. The protein was then mixed with the labeling (D_2_O) buffer, whose chemical composition was identical to the H_2_O crystallization buffer. The labeling reaction mixture consisted of 1 part protein (20 µM in H_2_O crystal buffer) and 9 parts D_2_O crystal buffer for a total D_2_O content of 90% for the reaction duration. The reaction was quenched after timepoints of 30 s, 100 s, 300 s, 1000 s, 3000 s, and 10,000 s. In order to stop the HDX labeling process, a cold quenching solution (1.5% formic acid, 3.0% acetonitrile, and 3 M GuHCl) was mixed in a 1:1 ratio with the labeled sample. The LEAP HDX system injected 100 µL of this solution into a pepsin column for digestion and further analysis. MS/MS fragmentation was used to confirm peptide identity along the sequence of PTP1B 1-321 WT, and these peptides and their retention times were used by the HDExaminer software to identify them automatically in the ’on-exchange’ experiments. In order to correct for back exchange in the regular and quantitative experiments, fully deuterated (FD) preparations were carried out on each PTP1B variant (PTP1B 1-321 WT, I19V, Q78R, D245G, P302Q). The protein was incubated at room temperature (20°C) for 28 days in a 90% D_2_O/H_2_O crystallization buffer. Duplicate (n=2) experiments were run for the 30 s and 300 s time points, while the 3000 s results are based on a single experiment in the WT, I19V, Q78R, and D245G variants.

### HDX-MS data analysis

#### Data preprocessing

Before performing the HDX analysis, the raw mass spectrometry data files (.d format) were processed using Compass Data Analysis 5.3 and Biotools 3.2 software to convert the data into a suitable format (.csv). The preprocessed data files were imported into version 3.3 of the HDExaminer software from Sierra Analytics. The imported data files were aligned based on the peptide identification information, retention time, and m/z values for accurate analysis. This step ensures that the corresponding peptide measurements from different time points are properly aligned for further analysis. The imported data were matched with the peptide sequences derived from the pepsin-based protein digestion within HDExaminer.

#### Peptide-level exchange rate analysis

The deuteration level of each peptide was determined by comparing the centroid mass of the deuterated peptide ion with that of the corresponding non-deuterated peptide ion at each time point. This calculation provides the deuteration percentage for each peptide at different time intervals. The HDExaminer software employs various algorithms to calculate the exchange rates of individual peptides. These algorithms utilize mathematical models, such as exponential fitting, to estimate the exchange kinetics and determine the exchange rates of the identified peptides. Relative deuterium uptake is expressed as the peptide mass increase divided by the number of peptide backbone amides. When the second residue of the peptide is not proline, the number of peptide backbone amides was decreased by one to account for rapid back-exchange by the amide adjacent to the N-terminal residue.

#### Visualization and data interpretation

The exchange rate profiles at the peptide and residue levels were computed in HDExaminer. The software provides graphical representations, such as heat maps and exchange rate plots, to facilitate the interpretation of the data. The exchange rate profiles can be further analyzed to identify protein regions exhibiting differential exchange behavior under different experimental conditions. To qualitatively visualize exchange values mapped to 3D protein structures, we obtained estimates of deconvoluted and smoothed residue-level interpolation of the peptide results from HDExaminer and plotted values along color spectra for Δ%deuteration (**Fig. 4**, **Fig. S7A**) that were then mapped onto the structures for the protein variants (I19V, Q78R, D245G, P302Q) relative to WT. For mapping Δ%deuteration for select peptides that span the sequence (**Fig. 4A**), we manually chose peptides that were moderate in length and efficiently spanned the sequence for ease of visual representation in the figure. For mapping Δ%deuteration for shorter-length peptides to the structure (**Fig. 5**), we prioritized shorter peptides to provide a closer comparison to the single-residue IADDAT data and still span the full sequence. HDX Δ%deuteration data for all peptides, mutants, and time points is available in **Table S7**.

### Structural modeling and visualization

The structural models for full-length PTP1B predicted by AlphaFold 2 (38) were obtained from the AlphaFold Protein Structure Database (39); these models include the catalytic domains plus the mostly disordered C-termini, which have not been structurally resolved experimentally, with varying annotated degrees of confidence for different residues. The sites of human variant mutations were mapped to the structures and visualized in the context of different ligands and crystal lattice contacts using PyMol (85).

Predictions of the effect of point mutations on structural stability were performed using FoldX v5 (40). This was done by first running the RepairPDB function of FoldX on four different structures of PTP1B including the closed and open states of RT apo PTP1B (6B8X), the WT structure (9CYO), and a full-length model from AF2. PositionScan was then used for each structure by specifying the four mutations of interest.

Final structural visualization and image production was performed using PyMol.

### Data availability

For X-ray crystallography, the crystal structure coordinates and structure factor data are available at the RCSB Protein Data Bank under the following PDB IDs (accession codes): 9CYO for WT, 9CYP for I19V, 9CYQ for Q78R, and 9CYR for D245G.

## Acknowledgements

We thank the physicians who referred people to the STILTS cohort and the participants for their involvement. We thank Alexander Wolff and Michael Thompson for IADDAT scripts and their assistance in running reciprocalspaceship, and Silvia Russi for assistance with RT data collection. The RT X-ray diffraction data reported here were collected at beamline 12-1 of the Stanford Synchrotron Radiation Lightsource (SSRL). Use of the SSRL, SLAC National Accelerator Laboratory, is supported by the U.S. Department of Energy, Office of Science, Office of Basic Energy Sciences under contract no. DE-AC02-76SF00515. The SSRL Structural Molecular Biology Program is supported by the DOE Office of Biological and Environmental Research and by the NIH, National Institute of General Medical Sciences (P30GM133894).

DAK is supported by NIH R35 GM133769 and a Cottrell Scholar Award. ISF is supported by a Wellcome Principal Research Fellowship (207462/Z/17/Z), National Institute for Health and Care Research (NIHR) Cambridge Biomedical Research Centre, Botnar Foundation, Leducq Foundation grant, Bernard Wolfe Health Neuroscience Endowment and a NIHR Senior Investigator Award. Part of this research has been conducted using the UK Biobank Resource under Application Number 53821.

