## Supplementary Materials for "Structures of human PTP1B variants reveal allosteric sites to target for weight loss therapy"

**The PDF file includes:**

Figures S1 to S8
Tables S1, S4, S5, S6

**Other Supplementary Materials for this manuscript include the following:**

Tables S2, S3, S7

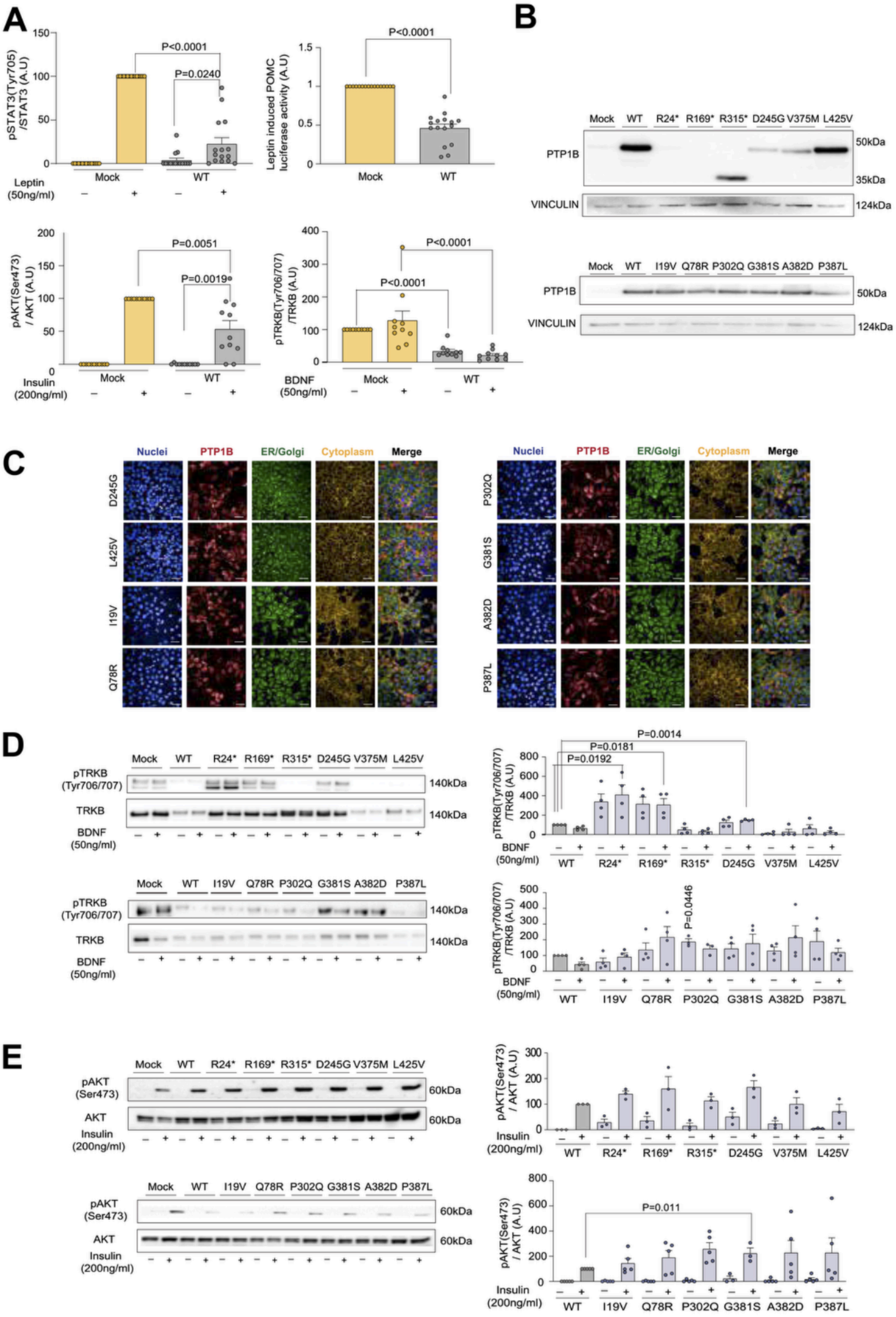


**Figure S1: Functional characterization of WT and mutant forms of PTP1B.**

(**A**) Effect of overexpression of human WT PTP1B in HEK293 cells on leptin-stimulated STAT3 phosphorylation (Tyr705) and POMC luciferase reporter activity, insulin-stimulated AKT phosphorylation

(Ser473) and BDNF-stimulated TRKB phosphorylation (Tyr706/707) compared to mock-transfected cells. n=10-17; mean +/- SEM, normalized to mock (0%) and mock stimulated (100%) (A.U: arbitrary units). Two-tailed unpaired one-sample t-test for WT PTP1B versus mock.

(**B**) Western blots depicting expression of WT/ mutant PTP1B in HEK293 cells, n=1.

(**C**) Representative confocal fluorescence microscopy images showing protein localization of WT/mutant PTP1B in HEK293 cells. Blue: DAPI for nuclei, red: Alexa 647 for HA tagged PTP1B, green: Alexa 488 for PDI, yellow: DyLight Phalloidin 554. Scale bar: 50 µm.

(**D**) Effect of WT/mutant PTP1B on BDNF-stimulated TRKB phosphorylation. n=3-4; data are expressed as mean +/- SEM normalized to WT (100%) (A.U: arbitrary units). Two-tailed unpaired one-sample t-test on log-transformed data for mutant versus WT.

(**E**) Effect of WT/mutant PTP1B on insulin-stimulated AKT phosphorylation. n=3-5; data expressed as mean +/- SEM, normalized to WT (0%) and WT stimulated (100%) (A.U: arbitrary units). Two-tailed unpaired one-sample t-test on log-transformed data for mutant versus WT.

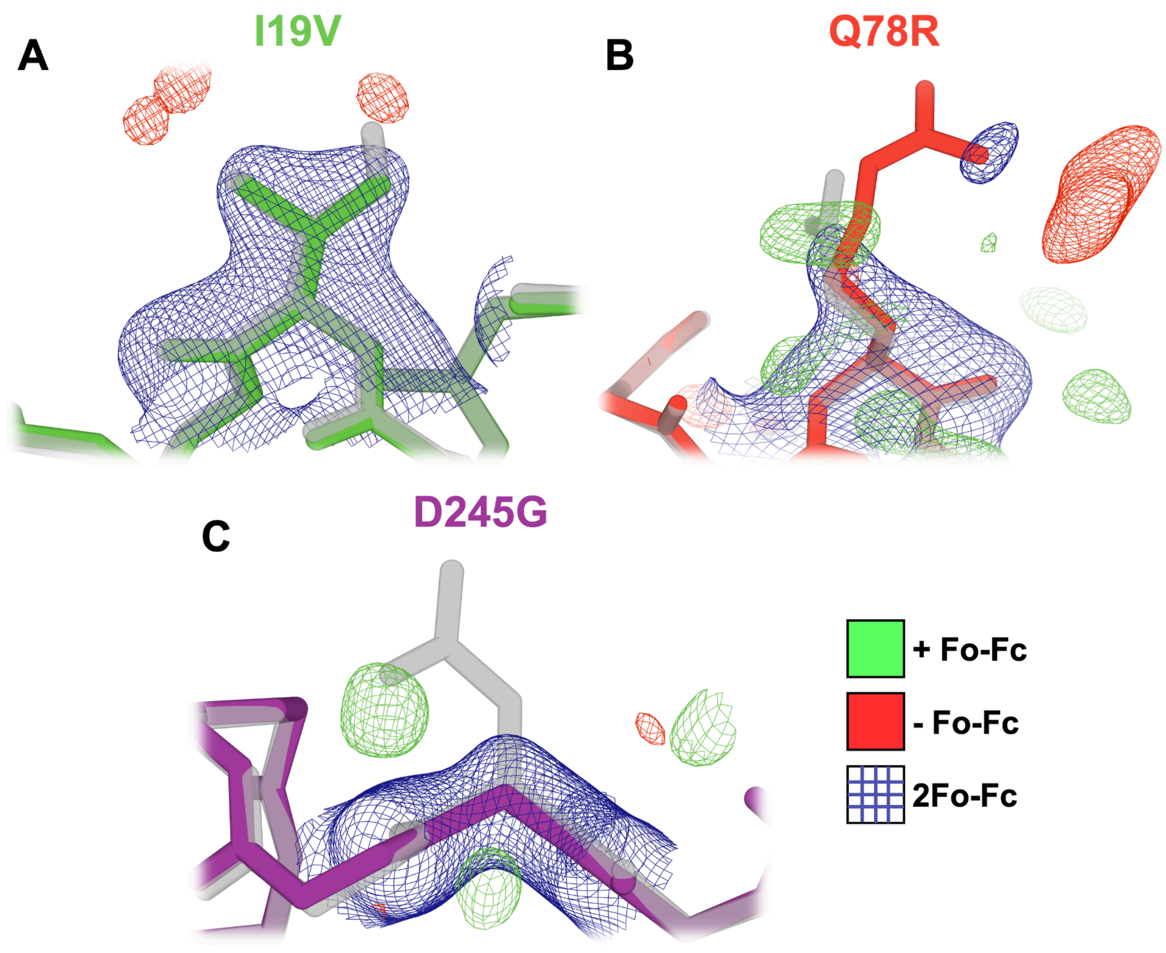


**Figure S2: Crystallographic electron density for mutations.**
Mutant crystal structure 2Fo-Fc and Fo-Fc electron density maps contoured at 1 σ for 2Fo-Fc (blue mesh) and +/- 3 σ for Fo-Fc (green/red mesh).
(**A**) The I19V mutant crystal structure (green sticks) presents no apparent 2Fo-Fc electron density for the delta carbon present in wild-type isoleucine (gray transparent sticks).
(**B**) The Q78R mutation site in its respective crystal structure (red sticks) is less clear. This is likely a function of the relatively poor resolution compared to the other mutant structures (2.30 Å in Q78R vs. 1.99 Å in I19V and 1.65 Å in D245G). Further, the exposure of the residue to solvent channels in the crystal makes it more likely for the side chain to adopt multiple conformations, and therefore challenging to resolve in electron density maps at low relative resolution.
(**C**) 2Fo-Fc electron density at the D245G mutation site (purple sticks) shows clearly that the entire wild-type aspartic acid side chain is absent from the D245G mutant crystal structure.

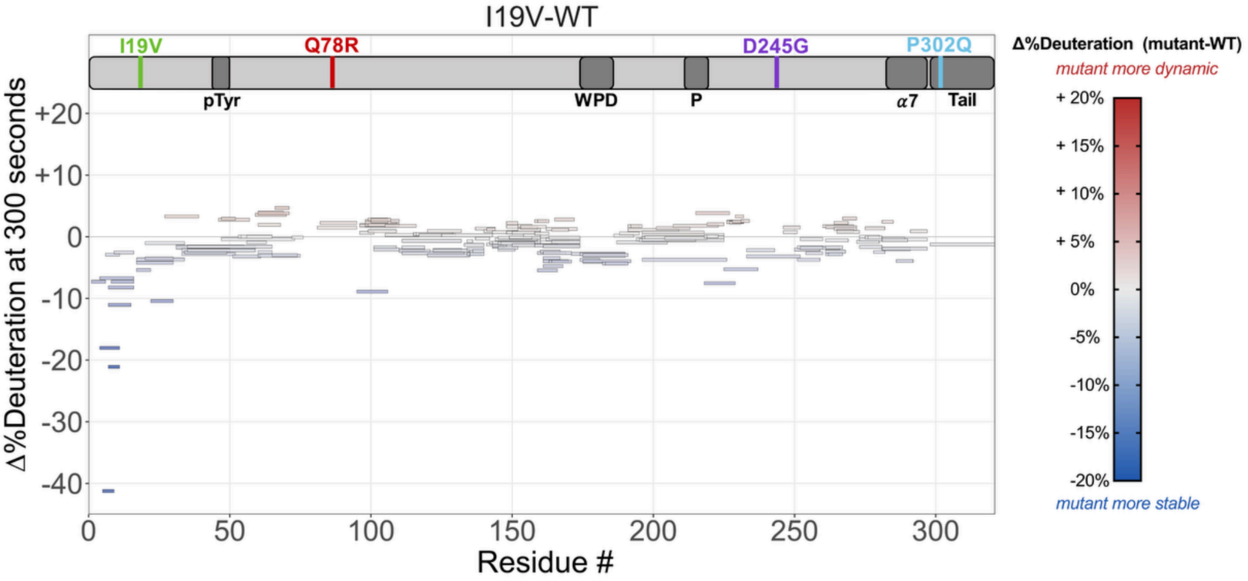


**Figure S3: HDX-MS difference Woods plots of I19V.**
The difference in %deuteration values at 300 seconds for the peptides of the I19V mutant PTP1B minus the values for WT PTP1B, plotted against amino acid sequence.

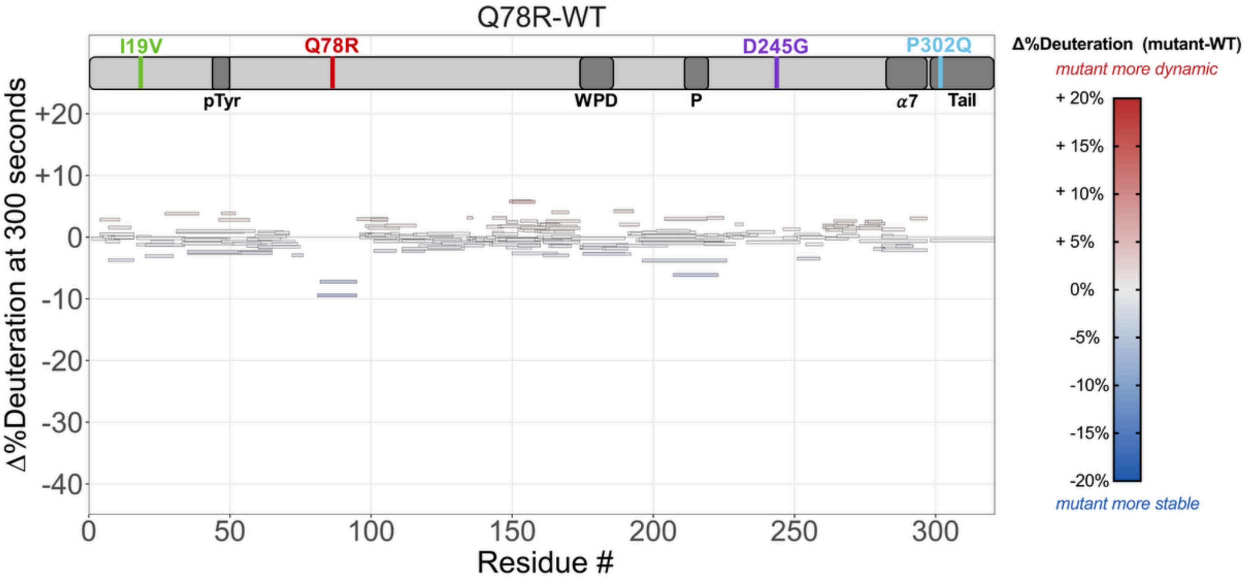


**Figure S4: HDX-MS difference Woods plots of Q78R.**
The difference in %deuteration values at 300 seconds for the peptides of the Q78R mutant PTP1B minus the values for WT PTP1B, plotted against amino acid sequence.

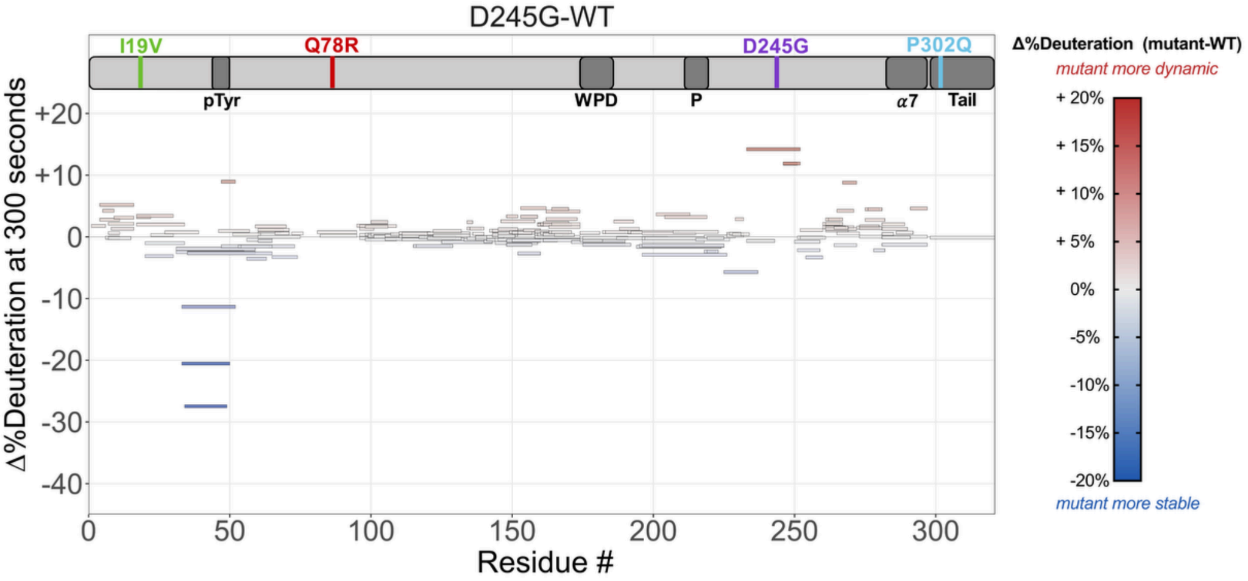


**Figure S5: HDX-MS difference Woods plots of D245G.**
The difference in %deuteration values at 300 seconds for the peptides of the D245G mutant PTP1B minus the values for WT PTP1B, plotted against amino acid sequence.

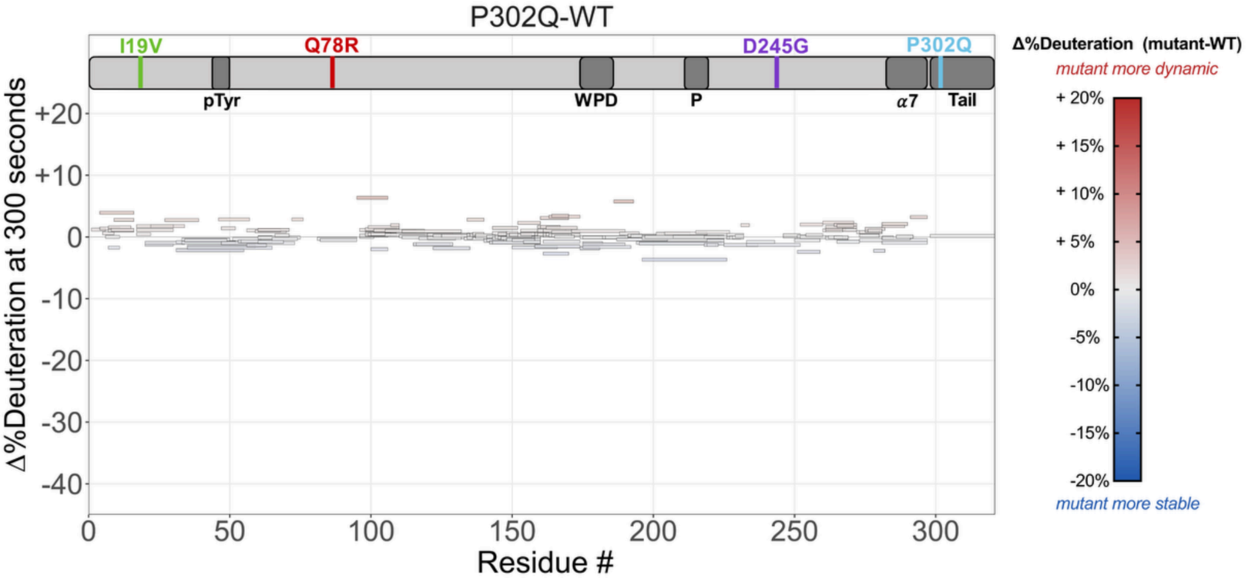


**Figure S6: HDX-MS difference Woods plots of P302Q.**
The difference in %deuteration values at 300 seconds for the peptides of the P302Q mutant PTP1B minus the values for WT PTP1B, plotted against amino acid sequence.

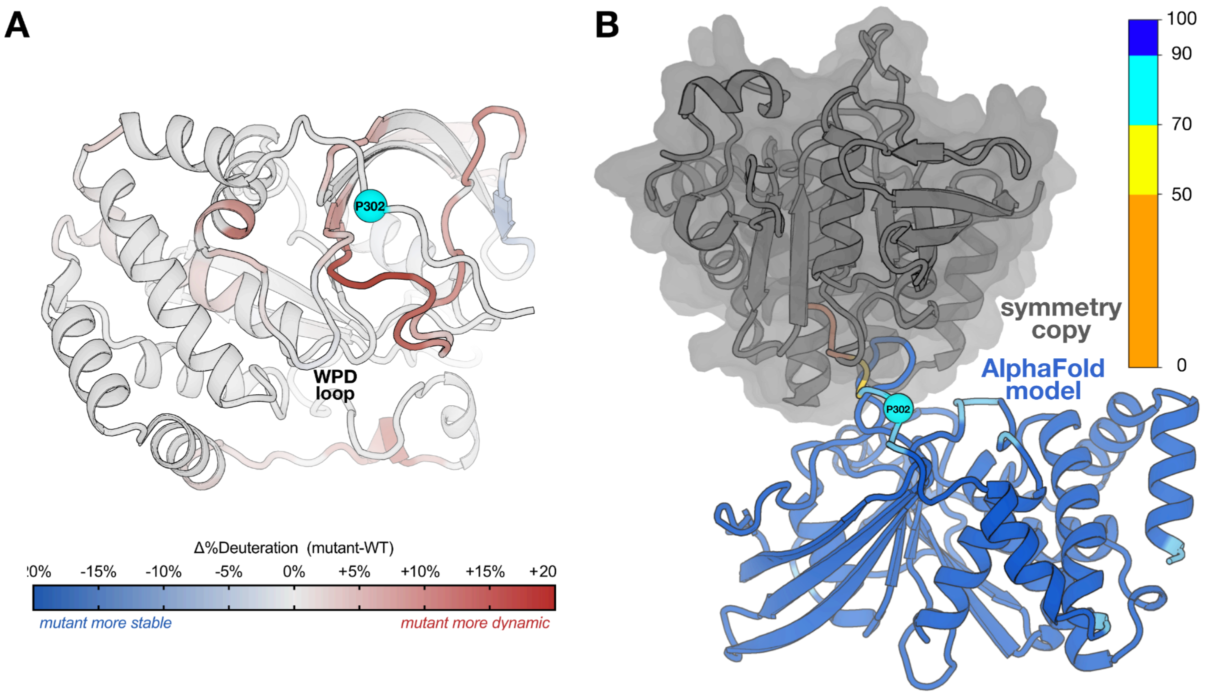


**Figure S7: Dynamic effects from a predicted quasi-ordered P302Q conformation.**
(**A**) HDX-MS results for P302Q-WT mapped onto a structural model obtained from AlphaFold DB (38, 39) only showing residues 1-310 for simplicity and to include the predicted location of the P302Q mutation. See color bar for corresponding mutant-WT difference HDX values at 300 seconds of labeling. Residues with Δ%deuteration between -5% and +5% are colored gray for visual clarity.
(**B**) The AlphaFold Database model (38, 39) of full-length PTP1B (residues 1-310 shown) colored according to the AlphaFold 2 predicted local distance difference test (pLDDT), along with an aligned symmetry mate from our $P3_121$ crystal form (gray), together show how crystal contacts impede the ability to resolve the predicted conformation of the C-terminal region including P302 in crystal structures.

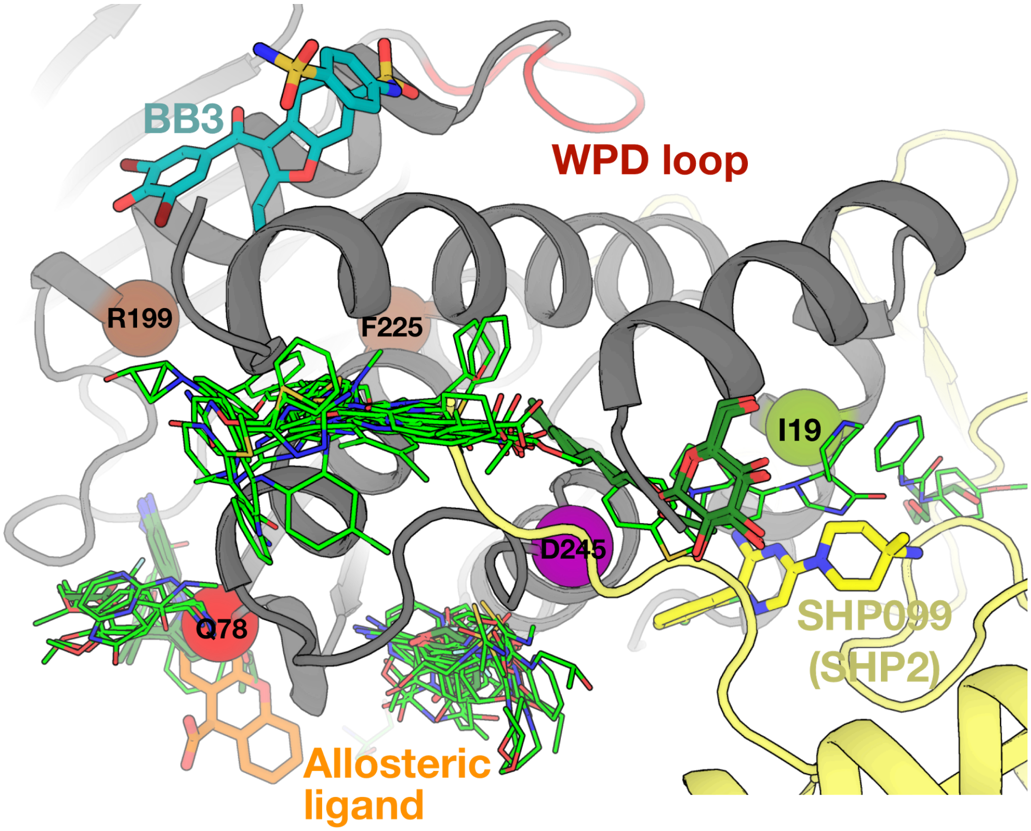


**Figure S8: SHP2 allosteric modulator binds near mutation sites.**
Same as **Fig. 5** with the addition of the allosteric modulator of the PTP1B homolog SHP2, SHP099 (yellow), and the SH2 domain of SHP2 (pale yellow) (PDB ID: 5EHR) (64).

| | Sequencing provider | | |
|---|---|---|---|
| | **BGI** | **Macrogen** | **Combined** |
| **Number of participants sequenced** | 399 | 598 | 997 |
| **Number of males (%)** | 64 (16) | 111 (18.6) | 175 (17.6) |
| **Number of females (%)** | 335 (84) | 487 (81.4) | 822 (82.4) |
| **Average age in years (SD)** | 43.2 (14.6) | 36.7 (13.5) | 39.3 (14.4) |
| **Average BMI in $kg/m^2$ (SD)** | 17.6 (0.93) | 17.5 (0.89) | 17.5 (0.91) |
| **Always thin (%)** | 356 (89.2) | 575 (96.2) | 931 (93.4) |
| **Family history of thinness (%)** | 301 (75.4) | 521 (87.1) | 822 (82.4) |

**Table S1: STILTS cohort participants who underwent whole exome sequencing.**

*Table S2 is provided as an XLSX file.*

**Table S2: PTP1B and TCPTP variants selected for molecular functional characterization from STILTS exomes and UK Biobank 200,000 exomes.**

*Table S3 is provided as an XLSX file.*

**Table S3: Statistical analysis of molecular assays for the characterization of PTP1B variants.**

| Variant | Location | Change in Function or Structure/Flexibility | | | |
|---|---|---|---|---|---|
| | | **Cell assays** | **Activity assays** | **X-ray structure** | **HDX-MS** |
| **I19V** | catalytic domain | no effect | significant catalysis decrease | effects in P loop and E loop | effects in α1’ helix |
| **Q78R** | catalytic domain | significant LOF | significant catalysis decrease | minimal effects | effects in P loop |
| **D245G** | catalytic domain | significant LOF | significant catalysis decrease | effects near active site and α4 helix | effects in pTyr loop |
| **P302Q** | Pro-rich region | significant LOF | no effect | --- | effects in α3 helix and 197 site |
| **V375M** | C-terminal tail | significant GOF; | --- | --- | --- |
| **G381S** | C-terminal tail | significant LOF | --- | --- | --- |
| **A382D** | C-terminal tail | significant LOF | --- | --- | --- |
| **P387L** | C-terminal tail | significant GOF | --- | --- | --- |
| **L425V** | ER anchor | no effect | --- | --- | --- |
| **R24*** | catalytic domain | significant LOF; | --- | --- | --- |
| **R169*** | catalytic domain | significant LOF | --- | --- | |
| **R315*** | Pro-rich region | no effect | --- | --- | --- |

**Table S4: Summary of functional and structural effects of PTP1B human variants.**

| | ΔΔG, *9CYO* (open) | ΔΔG, *6B8X alt A* (closed) | ΔΔG, *6B8X alt B* (open) | ΔΔG, AF2 (closed) |
|---|---|---|---|---|
| **I19V** | 0.66 | 1.51 | 0.35 | 0.45 |
| **Q78R** | -0.39 | 0.12 | -0.05 | -0.16 |
| **D245G** | 3.29 | 4.98 | 4.53 | 3.63 |
| **P302Q** | --- | --- | --- | 1.09 |

**Table S5: Predicted ΔΔG of mutations in/near catalytic domain using various PTP1B structures.**

FoldX was used to calculate the change in stability (folding free energy) for each variant: $\Delta\Delta G = \Delta G_{Mutant} - \Delta G_{WT}$ (kcal/mol). This was done for different models of PTP1B in either the "open" or "closed" state. Closed and open states were included as alternate conformations A and B in PDB ID 6B8X (previous structure of apo PTP1B at room temperature) and were extracted for separate analysis here. P302Q could only be modeled in the predicted AlphaFold 2 (AF2) model because residue 302 is disordered and absent from all existing crystal structures.

| PDB ID | *9CYO* | *9CYP* | *9CYQ* | *9CYR* |
|---|---|---|---|---|
| **Construct** | WT | I19V | Q78R | D245G |
| **Temperature (K)** | 298 K | 298 K | 298 K | 298 K |
| **Resolution (Å)** | 43.82–1.94 (2.01–1.94) | 44.76–1.99 (2.06–1.99) | 43.86–2.30 (2.39–2.30) | 38.79–1.65 (1.71–1.65) |
| **Completeness (%)** | 99.80 (99.92) | 99.78 (100) | 97.99 (98.40) | 99.88 (99.16) |
| **Multiplicity** | 9.7 (9.8) | 9.1 (9.0) | 10.2 (10.7) | 9.7 (7.2) |
| **I/sigma(I)** | 8.96 (0.83) | 6.02 (0.91) | 5.37 (0.37) | 16.67 (1.02) |
| **$R_{merge}$(I)** | 0.143 (2.884) | 0.187 (2.611) | 0.276 (3.299) | 0.053 (1.719) |
| **$R_{meas}$(I)** | 0.151 (3.043) | 0.198 (2.771) | 0.291 (3.466) | 0.056 (1.853) |
| **$R_{pim}$(I)** | 0.048 (0.961) | 0.065 (0.921) | 0.090 (1.055) | 0.018 (0.680) |
| **$CC_{1/2}$** | 0.928 (0.349) | 0.989 (0.354) | 0.987 (0.284) | 1.000 (0.417) |
| **Total observations** | 357343 (35614) | 311805 (30063) | 223386 (22975) | 580250 (42075) |
| **Unique observations** | 36906 (3639) | 34197 (3356) | 21897 (2151) | 59663 (5868) |
| **Space group** | P 31 2 1 | P 31 2 1 | P 31 2 1 | P 31 2 1 |
| **Unit cell dimensions (Å,Å,Å,°,°,°)** | 89.541, 89.541, 106.212, 90, 90, 120 | 89.518, 89.518, 106.110, 90, 90, 120 | 89.566, 89.566, 106.360, 90, 90, 120 | 89.579, 89.579, 106.234, 90, 90, 120 |
| **Solvent content (%)** | 45.85 | 43.33 | 42.93 | 48.34 |
| **$R_{work}$ (%)** | 16.95 | 18.20 | 16.23 | 16.65 |
| **$R_{free}$ (%)** | 20.15 | 21.39 | 20.73 | 18.58 |
| **RMS bonds (Å)** | 0.013 | 0.013 | 0.013 | 0.011 |
| **RMS angles (°)** | 1.226 | 1.250 | 1.230 | 1.124 |
| **Ramachandran outliers (%)** | 0.36 | 0.35 | 0.35 | 0.35 |
| **Ramachandran favored (%)** | 96.80 | 96.45 | 96.10 | 97.16 |
| **Clashscore** | 1.06 | 2.33 | 2.13 | 2.08 |

**Table S6: Crystallographic statistics.**
Overall statistics given first (statistics for highest-resolution bin given in parentheses).

*Table S7 is provided as an XLSX file.*

**Table S7: HDX-MS values for all catalytic domain PTP1B variants.**

Change in HDX rate (mutant-WT ∆%deuteration) for HDX-reportable residues between mutant and wild-type for each peptide.
